# Systematic evaluation of an exhaustive set of connectivity estimators in bivariate and multivariate modes for an improved virtual source MEG connectivity analysis for the study of resting state recordings

**DOI:** 10.64898/2026.02.21.707012

**Authors:** SI Dimitriadis

**Affiliations:** Integrative Neuroimaging Lab, 55133, Thessaloniki, Makedonia, Greece; 1^st^ Department of Neurology, G.H. ‘AHEPA’, School of Medicine, Faculty of Health Sciences, Aristotle University of Thessaloniki (AUTH), 54124, Thessaloniki, Makedonia, Greece; Neuroinformatics Group, Cardiff University Brain Research Imaging Centre (CUBRIC), School of Psychology, College of Biomedical and Life Sciences, Cardiff University, Maindy Rd, CF24 4HQ, Cardiff, Wales, United Kingdom; Neuroscience and Mental Health Research Institute, Cardiff University, Cardiff, CF24 4HQ, UK; MRC Centre for Neuropsychiatric Genetics and Genomics, Division of Psychological Medicine and Clinical Neurosciences, Cardiff School of Medicine, Cardiff University, Cardiff, CF24 4HQ, UK

**Keywords:** Magnetoencephalography (MEG), Functional Connectivity, Virtual Source Space, Multivariate Connectivity, Brain Connectomics, Biomarkers

## Abstract

Brain activity is measured using noninvasive electrophysiological techniques, such as electroencephalography (EEG) and magnetoencephalography (M/EEG), recorded from sensors outside the skull and it is regularly transformed into a virtual source space which is parcellated into anatomical brain areas using an atlas. Then, functional connectivity (FC) is estimated between pairs of regions, with their brain activity characterized by a representative time series extracted from multiple voxel time series (multidimensional), using various techniques. Alternatively to bivariate FC estimators and these techniques, their multivariate representative-free extension has been proposed. Multivariate FC estimators can also be considered. The impact of these different strategies on FC estimation is still unclear.

An appropriate framework for systematically evaluating FC estimators in the virtual MEG space and across multiple processing steps for brain network construction is missing. Here, we compared an exhaustive set of bivariate FC estimators paired with techniques for extracting representative time series, their multivariate extensions, and multivariate estimators under two scenarios: 1) on the discrimination of MCI subjects versus healthy controls, using a k-NN classifier and an appropriate graph distance metric, and 2) on the repeatability of brain network topologies in a test-retest study.

Our results demonstrate that the multivariate extension of bivariate FC estimators (representative-free approach), which summarizes pairwise FC strength across all voxels of two brain areas, and accurate multivariate estimators that consider pairs of region-wise voxel time series at once, clearly outperform bivariate FC estimators paired with representative time series in both scenarios. Given the remarkable differences across the three approaches consistently across the frequency bands and FC estimators, caution is warranted when comparing results across studies that apply different pairs of extraction methods and FC estimators.

## 1. Introduction

Magnetoencephalography (MEG) and electroencephalography (EEG) are widely used non-invasive techniques for measuring electrophysiological brain activity [1–3]. Both neuroimaging modalities have lower spatial resolution than functional magnetic resonance imaging [4]. This significant drawback of MEG and EEG poses challenges for accurate neural source localization. This difficulty stems from the inherently ill-posed nature of the inverse problem in source localization: the number of potential neural source locations far exceeds the number of scalp sensors, yielding an infinite set of possible source configurations that could explain the recorded signals [5]. However, they are known for their high millisecond-scale temporal resolution [3] and provide key insights into the intricate dynamics of healthy [6], neurological [7–9], and neuropsychiatric [10] clinical populations.

The transition from recording scalp sensors to brain virtual source generators requires adopting a source reconstruction approach [10]. Inverse source localization solutions can be broadly classified into two main categories (Grech et al., 2008; Baillet et al., 2001): those focusing on distributed source imaging and those on dipole fitting. Distributed source imaging models hypothesize that virtual brain sources are distributed across the cortical surface, and source localization algorithms estimate a density map of these sources using linear optimization methods such as the minimum-norm estimate (MNE) [3,5], weighted MNE [11,12], low-resolution electromagnetic tomography (LORETA) [13], standardized LORETA (sLORETA) [14], etc. However, these methods typically yield spatially diffuse solutions that blur focal sources due to the ill-posedness of the inverse problem and the effects of regularization constraints.

In contrast, dipole fitting methods avoid the ill-posedness of the inverse problem by identifying a small set of equivalent current dipoles (ECDs) whose electromagnetic fields best match the M/EEG measurements using a least-squares approach [15]. Rather than assigning a current value to every possible dipole in a densely sampled source space, these methods focus on identifying specific focal sources. Key dipole fitting methods include beamformers [16,17] and MUSIC [15], as well as recursive extensions such as RAP-MUSIC [18], Truncated RAP-MUSIC [19], and RAP Beamformer [20]. A recent study introduces PATCH-AP, an enhanced version of the Alternating Projection (AP) method that effectively localizes both discrete and spatially extended sources. This new source localization approach addresses the limitations of both categories of electromagnetic source localization methods [21].

The initial mathematical description of inverse models was based on the hypothesis that only a small number of sources, termed dipoles, were active at any given time [22]. In the last two decades, we have used more complex models that assume multiple brain virtual sources are simultaneously active, producing the electromagnetic activity measured on the scalp [5,16]. The source models range from a few thousand (spatial filter approaches) to tens of thousands (distributed source models) sources. The EEG and MEG sensor layout consists of tens to a few hundred sensors.

The EEG and MEG sensor arrays range from tens to a few hundred sensors [23], and the resulting source-space brain activity is distributed across thousands of voxel time series, which is highly rank-deficient. Estimating FC using rank-deficient data yields many ‘ghost’ interactions [24], diminishing the usefulness of this approach. Moreover, using voxel time series as input to FC estimations yields large, redundant whole-brain FC matrices. In contrast, researchers use a brain atlas with a limited number of brain areas, comparable to the number of sensors, to estimate FC between these areas [25–27]. This approach minimizes the effect of the matrix’s redundancy on FC estimations.

Each brain atlas summarizes voxel time series into groups aligned with the spatial boundaries of a brain area. To characterize the brain activity of a given area from a set of voxel time series that encapsulate the convex hull of that area, we need to apply one of the available methods. Several approaches have been proposed to define representative time series for each brain area, thereby characterizing its activity. Most recent studies use techniques based on representative time series, in which a single time series captures the entire activity of each area (e.g., identification of the center of mass (CENTROID), time series with maximal power (POWER), weighted average (INTERPOLATION), etc.). [28–33].

However, these approaches oversimplify the representation of brain-area-based activity, leading to information loss. To overcome this limitation, many researchers have developed fully multivariate extensions of well-known FC metrics that integrate multidimensional information from both brain regions into a single FC estimate [34–39].

A ‘multivariate’ strategy acknowledges that all elements of the source space and their time courses carry some useful information. At the same time, the aggregation approach, such as mean or PCA, is the easiest computational choice. The use of these multivariate approaches has, for example, been proven to improve the accuracy of the area-to-area FC estimation in functional magnetic resonance imaging [40]. Basti and colleagues provided a comprehensive review of these methods, along with a small toolbox of examples [41]. In another study, the authors proposed a method to estimate representative free-phase synchronization by extending bivariate phase estimators to the multivariate case [38]. They demonstrated the superiority of this approach over well-known methods for estimating representative time series for each brain area in simulations involving targeted pairs of virtual brain sources. Complementary, they showed in an EEG-MEG virtual source connectivity study a high correlation between the FC values estimated from EEG and MEG, suggesting that the results found in the high-sensitivity, low-noise MEG can be transferable to the more affordable EEG modality [39]. Brkic et al. [37] performed an analysis using a large resting state MEG cohort (N=113) with four extraction methods (mean before, PCA before, mean after, and max after) and two connectivity measures (ciPLV, PLV), across three frequency bands, by applying one canonical cortical atlas (Desikan). They concluded that, after dimensionality reduction in voxel-by-voxel FC analysis, it should be preferred when higher sensitivity is desired, despite the higher computational demands.

An appropriate framework for systematically evaluating FC estimators in the virtual MEG space and across multiple processing steps for brain network construction is missing. In this work, we compared an exhaustive set of bivariate FC estimators (240) with a combination of techniques for extracting representative time series per ROI (6), their multivariate extensions by aggregating the voxel-by-voxel FC (3), and multivariate estimators that simultaneously consider pairs of voxel time series between every pair of ROI (7). Our question was how the choice of ROI extraction method, their multivariate extension, and original multivariate FC estimators affect the estimation of MEG resting-state functional connectivity; Our question was addressed under two scenarios: 1) the discrimination of healthy controls and MCI subjects across seven frequency bands using a k-NN classifier and an appropriate graph distance metric, and 2) the repeatability of functional connectivity brain networks (FCBN) topologies across seven frequency bands in test-retest study.

We hypothesized that both the multivariate extension of bivariate estimators to a representative-free approach and the multivariate estimators that simultaneously summarize brain synchronization between a pair of brain areas across multiple voxel time series would have outperformed those based on representative time series and bivariate FC estimators. In short, this work addresses the following questions : (i) How does the ROI extraction method affect the estimation of FCBN at resting-state reflected in the discrimination accuracy of MCI vs healthy controls, and in the repeatability of FCBN ?, (ii) Are differences over discrimination accuracy and repeatability of FBN topologies between extraction strategies (representative time series per ROI vs multivariate extensions of bivariate FC estimators vs multivariate FC estimators) consistent across an exhaustive set of FC estimators and seven frequency bands ?, and (iii) What are the recommendations for future studies at MEG resting-state functional connectivity ?.

## 2. Methods

### 2.1 Approaches based on representative time series

The most common approach to performing FC analysis in an atlas-based, parcellated virtual-source brain space is to assign a representative time series to define the overall activity of a brain area. These representative time series can be linked to an existing voxel time series within the convex hull of a brain area or to a transformation of voxel time series corresponding to different virtual source positions within the spatial limits of a brain area.

In the first category, the most common approach is the use of the virtual source closest to the brain area centroid (CENT) or the use of the virtual source associated with the most significant power (POWER) [42]. The POWER method is more appropriate for targeting brain areas such as the visual area than whole-brain approaches, which can introduce depth biases due to deep sources in beamformers [43]. In another study, the authors attempted to detect the virtual source by identifying the time series most correlated with the remaining virtual sources within the brain area [32].

In the second category, the goal is to combine all the virtual source time series anatomically constrained by the adopted atlas within a brain area to extract the representative time series. A common practice in functional magnetic resonance imaging (fMRI) FC analysis is to average the time series of all sources within a brain area to obtain a representative time series (AVERAGE) [44]. A more popular choice is principal component analysis (PCA), which weights the virtual source activities and combines them so that the principal components capture the maximum possible variance across the whole brain [33,45]. We also proposed an interpolation approach that weights each virtual source time series according to its correlation with the remaining virtual source time series within the brain area (INTERPOLATION) [6,9,46]. A fourth way to combine all the brain area’s virtual source time series into a representative time series is to employ the Kosambi-Hilbert torsion (KHT) [47]. This method extends the PCA-based optimization approach to extract the dominant activity in noisy environments. As the source-reconstructed activity is a noisy estimate of the brain area’s oscillatory activity, KHT extracts the oscillatory brain activity with the highest signal-to-noise ratio. KHT is not a high-frequency method in FC analysis, but it was used in a simulation study [38]; therefore, I included it in the overall analysis.

In this study, we will evaluate the performance of six methods based on representative time series: two from the first category (CENT, POWER) and four from the second category (AVERAGE, PCA, INTERPOLATION, KHT). Their performance will be evaluated alongside a comprehensive set of FC estimators, as described in the following sections. These approaches reduce the description of the brain area’s activity from multi-voxel time series to a single time series and the FC to (1D) one-dimensional connectivity. However, both categories of representative approaches ignore potential (MD) multi-dimensional connectivity between two regions. Below, I will describe and employ two techniques to overcome this drawback. The first involves estimating synchronization between the time series of every pair of brain areas (Section 2.2), and the second uses multidimensional connectivity estimators that operate on the two sets of voxel time series (Section 2.4).

### 2.2 Multivariate extension of bivariate FC estimators

Bivariate FC estimators that quantify synchronization between pairs of brain areas can be extended to their multivariate counterparts by constructing a pairwise synchronization matrix. Suppose one area contains N virtual sources and another one M virtual sources. In that case, all pairwise associations between two brain areas can be represented by a non-symmetric N × M matrix that tabulates the bivariate FC estimates. The authors who proposed the Phase Slope Index [48] extended it to a multivariate counterpart [36] by employing operations on this matrix.

A straightforward approach to account for all possible interactions between the two areas is to obtain a single value equal to the average of the N x M matrix. This global value is the average of the entries in the bivariate FC matrices [49,50]. I will term this approach FC-AVG. Similarly, we will use the median of the N × M matrix; this approach will be referred to as FC-MEDIAN hereafter.

Previous studies described how to obtain a single value from a rectangular matrix that tabulates all pairwise synchronization strengths between L sources of size L x L. A common approach was to extract this value via clustering analysis of the matrix’s eigenanalysis. The eigenvectors can inform us about the cluster membership of every source L, while the eigenvalues indicate the global variance explained by the cluster [51,52]. This approach can be used to define the global functional strength of this matrix as the summation of the normalized eigenvalues [38].

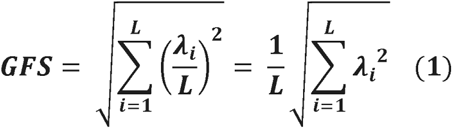

In (1), λ_i_ is the i-th eigenvalue of the pairwise Functional Strength (FS) matrix.

An alternative, equivalent way to integrate the strengths of all pairwise synchronizations tabulated in an N x M matrix is to use the normalized Frobenius norm. The Frobenius norm can be calculated for rectangular nonsymmetric matrices, and an index of inter-area FC estimations can be obtained using bivariate FC estimators [38]. We call it hereafter Global Functional Strength (GFS), the global measure of the functional strength of this N x M matrix, and it is expressed as:

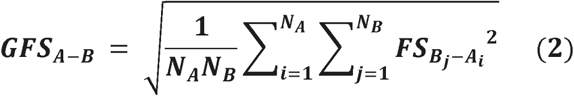

In (2), N_A_ is the number of sources in area A, and Ai is the i-th source of area A. GFS is equal to the root-mean-square (RMS) of all the elements in the bivariate FS matrix. We will term this approach, hereafter as FC-RMS.

In this work, we will evaluate the performance of three multivariate methods: the average of all bivariate FC values (FC-AVG), the median of all bivariate FC values (FC-MEDIAN), and the root mean square of all bivariate FC values (FC-RMS).

### 2.3 An exhaustive set of bivariate FC estimators

We used the recently developed Python Toolkit for Statistics on Pairwise Interactions (pyspi; v0.4.1, commit c19d06) to calculate alternative measures of pairwise interactions (SPIs) for FC. A recent study introduced an extensive, assembled library of 237 statistics for pairwise interactions [53]. Recently, researchers estimated pairwise FC associations between pairs of brain areas using this Python module in resting-state functional magnetic resonance imaging (fMRI) datasets [54]. In addition to this large set of statistics for pairwise interactions, I added the imaginary part of the phase locking value (iPLV) [6,9,33,44], the corrected imaginary part of the phase locking value (ciPLV) [55], and the Circular-circular correlation (CCc) [56]. The total number of pairwise FC estimators was 240.

### 2.4 Multivariate FC estimators

However, the aforementioned summary approach ignores potential multivariate (MV) connectivity between two regions, and many recent methods have been proposed to capture such complex dependencies.

We then consider five different, time-domain MV-connectivity methods: “canonical correlation” [34], “multivariate pattern dependence” [57], “distance correlation” introduced by [58] and applied for FC [40,59] and structural connectivity [40], “representational connectivity analysis” [60], and “linearly predicted representational dissimilarity” [61]. These represent the prototypical cases of all the primary methods that we are aware of, which have been proposed and are currently used in fMRI. For the frequency domain, we focus on phase-coupling methods. In particular, I employed the MV versions of two 1D phase-coupling methods, i.e., “imaginary coherency” [62] and “lagged coherence” [63]. These MV generalizations are termed the “multivariate interaction measure” [35] and the “multivariate lagged coherence” [64]. These are the primary MV phase-coupling methods we are aware of. A total of seven MV FC estimators will be tested in the present study.

### 2.5 Methodology and Hypotheses

Our study first evaluated the superiority of using all available information within each brain region for FC estimation over classical approaches that rely on a single representative time series. At the second level, we compared multivariate FC estimators with bivariate FC estimators that use all information within each brain region. We hypothesized that multivariate FC estimators would outperform the multivariate extension of bivariate FC estimation, as well as classical approaches based on a representative time series.

## 3. Description of the MEG resting-state datasets

### 3.1 Connectomic Whole-Brain Biomarkers for MCI vs HC as a common comparative framework

#### 3.1.1 Description of the MEG sample in brief

Data were obtained from 24 subjects diagnosed with mild cognitive impairment (MCI) (11 males, mean ± SD age 72.77 ± 3.31 years) and 30 healthy controls (13 males, mean ± SD age 72.37 ± 2.63 years). The MCI and control groups were recruited from the Hospital Clínico Universitario San Carlos (Madrid). All subjects were right-handed and native Spanish speakers [65]. Table 1 summarizes the demographic characteristics and mean hippocampal volumes for subjects in both groups.

**Table 1.** Mean ± standard deviation of the demographic characteristics of controls and MCIs.

|  | Number of subjects | Gender (M/F) | Age | MMSE | LH ICV | RH ICV |
| --- | --- | --- | --- | --- | --- | --- |
| Control | 30 | 13/17 | $72.37 \pm 2.63$ | $29.13 \pm 0.94$ | $0.0026 \pm 0.0003$ | $0.0026 \pm 0.0003$ |
| MCI | 24 | 12/11 | $72.67 \pm 3.31$ | $26.43 \pm 3.22$ | $0.0021 \pm 0.0003$ | $0.0022 \pm 0.0004$ |

*M, males; F, females; MMSE, Mini-Mental State Examination; LH ICV, Left hippocampus normalized by total intracranial volume (ICV); RH ICV, Right hippocampus normalized by ICV.*

Data acquisition parameters for structural and functional neuroimaging, the denoising strategy for MEG recordings, and the source localization approach are described in our previous study [33]. Below, we provide links to download the open MEG-MCI dataset.

1. Dataset part I (Controls):https://figshare.com/s/71a5fb9043235740a6a7 doi: 10.6084/m9.figshare.6210158
2. Dataset part II (MCI):https://figshare.com/s/9660b976e4138853d845 doi: 10.6084/m9.figshare.5858436

We selected, for each subject, multiple artifact-free trials of 6 s (6,000 samples) after careful visual inspection, yielding 32–44 epochs for further analysis. Time-series of neuronal activation were computed for the seven frequency bands: δ (0.5–4 Hz), θ (4–8 Hz), α_1_ (8–10 Hz), α_2_ (10–13 Hz), β_1_ (13–20 Hz), β_2_ (20–30), γ_1_ (30–45 Hz) using a third order Butterworth filter with zero-phase using filtfilt.m function from MATLAB.

#### 3.1.2 Functional Connectivity Brain Network Topologies

We first concatenated multiple artifact-free 6s trials at the voxel level. Then, we extracted frequency-dependent voxel-wise resting-state recordings using Linearly Constrained Minimum Variance (LCMV) [16]. The virtual space is parcellated using the AAL (Automated Anatomical Labeling) atlas, which groups voxel time series into 90 brain regions, 45 per hemisphere. Subsequently, we estimated functional connectivity brain networks for each computational scenario, which are expressed as 2D tensors (matrices) of size [90 x 90]. In detail, we computed across the seven frequency bands individualized functional connectivity brain networks :

***1)*** (representative time series) for the combination of 6 approaches for extracting the representative time series per brain region multiplied by the 240 connectivity estimators (6 x 240)
***2)*** (multivariate extension of bivariate FC estimators) for the combination of 3 approaches for extracting the summary of FC metric value working on the voxel level (representative-free approach) multiplied by the 240 connectivity estimators (3 x 240)
***3)*** (multivariate estimators) for the adaptation of seven (7) multivariate FC estimators that simultaneously manage the multi-voxel time series within every pair of brain areas

Fig.1 outlines the preprocessing steps from the MEG sensor activity to the construction of functional brain networks (FBN).

**Fig. 1.**
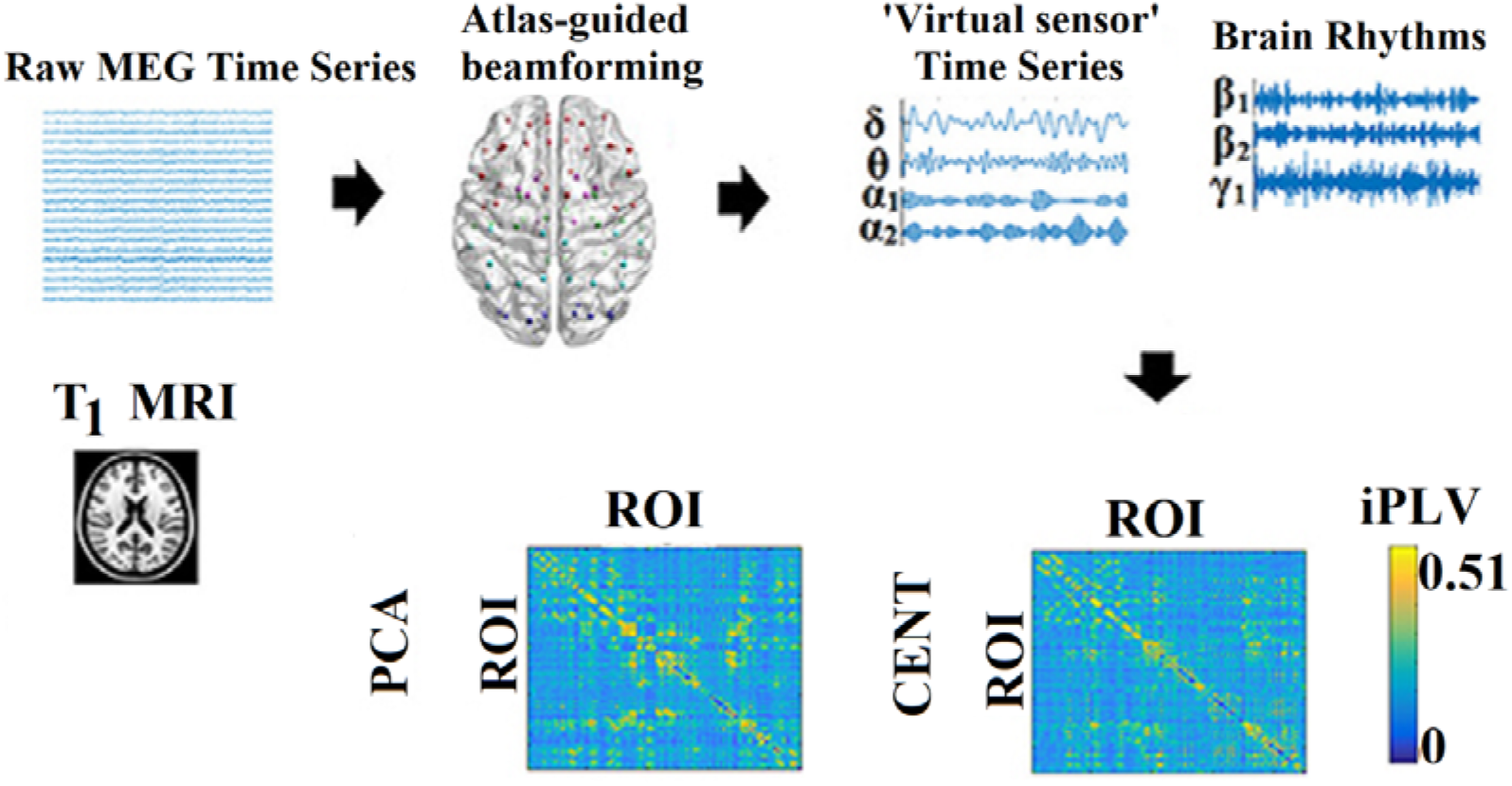
A flowchart of the processing steps from the MEG sensor level recodings to the construction of the FBN on the virtual source space.

#### 3.1.3 Data-driven Topological Filtering of Functional Brain Networks with OMST

Each individualized FBN was topologically filtered using our data-driven OMST approach [66,67]. OMST was applied across frequency bands, estimators, and methods. For each FBN, we estimated sparsity after applying OMST to the fully weighted network. Sparsity is measured as the ratio of the number of selected functional edges to the total number of possible pairs of brain regions; here, 90*89/2 = 4.005. Every OMST captures 89 functional edges (90 - 1 = 89), corresponding to a sparsity level of 89/4005 = 2.22%. The derived estimated sparsity levels will be equal to multiples of 2.22%.

#### 3.1.4 Classification Approach

We evaluated our methodological strategies across three directions of FC estimation in the virtual source space by comparing the classification performance of FBN topologies between MCI and control groups. Classification performance of the derived FC brain network topologies was realized following a 5-fold cross-validation and by adapting a k-NN classifier (with k = 5) using portrait divergence (PID) as a proper graph distance metric [44,68]. Classification accuracy, sensitivity, and specificity were estimated per case.

Fig.2 illustrates how the adopted kNN-PID classification scheme operates on the estimated FBNs from the two subject groups.

**Fig. 2.**
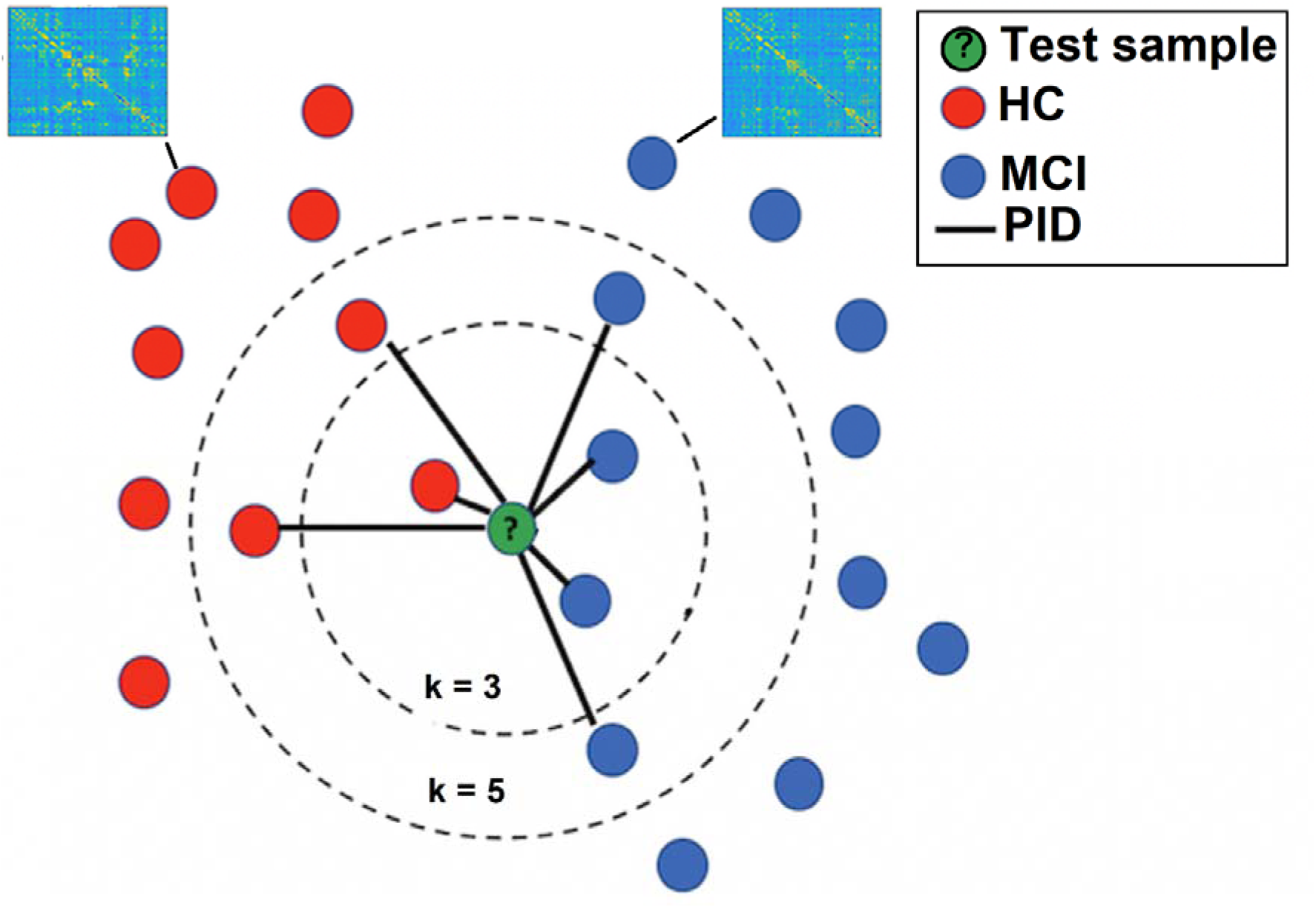
A visualization of a k-Nearest Neighbor (kNN) classifier tailored to the FBN topologies.

### 3.2 Repeatability of MEG resting-state FCBN in a test-retest study

#### 3.2.1 Human Connectome Project (HCP) data

The dataset for this study comprised structural and functional data, specifically resting-state magnetoencephalography (MEG) and functional magnetic resonance imaging (fMRI), from 89 subjects (46% female, mean age 29.0 ± 3.6 years) selected from the 1200 Subjects release of the Human Connectome Project (HCP) [69–71]. All participants included in the analysis had complete anatomical, resting-state MEG, and fMRI data, and provided written informed consent in accordance with HCP consortium guidelines. Resting-state MEG recordings were obtained at St. Louis University using a whole-head MAGNES 3600 system (4D Neuroimaging, San Diego, CA) equipped with 248 magnetometers and 23 reference channels. Data were sampled at 2034 Hz in three separate runs, each lasting approximately six minutes, during a single-day recording session. During acquisition, subjects were positioned supine in the scanner with eyes open. Electrooculography (EOG) and electrocardiography (ECG) were recorded to facilitate rejection of ocular and cardiac artefacts. Additionally, the outline of each subject’s scalp (approximately 2400 points), anatomical landmarks, and localizer coil positions were digitized at the start of the recording session. A structural T1w volume with 0.7 mm isotropic voxel size was acquired as well. Our analysis focused on sessions 1,2 (temporally closed sessions), and session 3 vs 1,2 (temporally distant sessions).

#### 3.2.2 MEG Processing

We downloaded the preprocessed sensor-level MEG data from the HCP database. The MEG preprocessing pipeline includes the following steps: (1) Rejection of bad channels: we removed non-working channels, (2) Filtering: band-pass filtering (1–150 Hz) and notch filtering (59–61 Hz/119–121 Hz) to remove power line artefacts, and (3) Denoising of MEG multi-channel recordings with wavelet ICA (wICA): As in our previous studies [59,72,73], we adopted wICA as a way to 1) first decompose MEG multi-channel recordings into N ICs, where N denotes the number of MEG sensors.

The time course of each independent component (IC) was decomposed using Daubechies wavelet filters. Wavelet decomposition of IC broadband activity was performed on approximately 180 2-seconds temporal segments from 6-minute recordings in every session. Each wavelet subcomponent of an IC was classified as either an ocular, muscle, or cardiac artifact, or as genuine brain activity, based on visual inspection and supported by kurtosis, skewness, and entropy measures [59,72,73]. For each 2-seconds segment, artifactual wavelet segments were set to zero, and the cleaned 2-seconds IC was reconstructed. Subsequently, the cleaned 2-seconds ICs were recomposed into cleaned IC time courses, which were further recomposed into cleaned wavelet-ICA MEG activity for each temporal segment. In this study, the total number of wavelet subcomponents was not completely rejected in any temporal segment across ICs and subjects. Power spectrum analysis was performed for each subject before and after the correction procedure, resulting in more pronounced characteristic frequency peaks. The wavelet plus ICA (wICA) method offers the advantage of correcting artefactual ICs across the time course, in contrast to zeroing the entire IC, which would also eliminate genuine brain activity.

#### 3.2.3 Cortical Parcellation

After artefact correction of MEG resting-state with wICA, we source-localized MEG activity. We extracted frequency-dependent voxel-wise resting-state recordings using Linearly Constrained Minimum Variance (LCMV) [16]. The virtual space is parcellated using the AAL (Automated Anatomical Labeling) atlas, which groups voxel time series into 90 brain regions, 45 per hemisphere. As in the first dataset, we computed across the seven frequency bands individualized functional connectivity brain networks :

1. (representative time series) for the combination of 6 approaches for extracting the representative time series per brain region multiplied by the 240 connectivity estimators (6 x 240)
2. (multivariate extension of bivariate FC estimators) for the combination of 3 approaches for extracting the summary of FC metric value working on the voxel level (representative-free approach) multiplied by the 240 connectivity estimators (3 x 240)
3. (multivariate estimators) for the adaptation of seven (7) multivariate FC estimators that simultaneously manage the multi-voxel time series within every pair of brain areas

#### 3.2.4 Data-driven Topological Filtering of Functional Brain Networks with OMST

Each individualized FBN was topologically filtered using our data-driven OMST approach [66,67]. OMST was applied across frequency bands, FC estimators, methods, and sessions.

#### 3.2.5 Repeatability of FCBN between sessions

We evaluated our methodological strategies across three directions of FC estimation in the virtual source space by quantifying the graph distance between individual FCBN related to the three sessions. For that purpose, we adopted “Portrait divergence” (PDiv), a measure of dissimilarity between networks [68], as we did in a previous study with fMRI modality [44]. PDiV quantified the dissimilarity between FCBN linked to sessions in a pair-wise fashion (session 1 - session 2, session 1 - session 3, session 2 - session 3). We finally estimated the mean PDiV between the three pairs per individual across frequency bands, FC estimators, and methods.

### 3.3 Statistical Analysis and Post hoc tests

Statistical and post hoc tests were performed to compare the performance of the classification approach across the methodological approaches (six for representative time series, and three for the multivariate extension of bivariate estimators) independently for each frequency band. For the first dataset, we compared classification performance, sensitivity, and specificity. For the second dataset, we compared the PDiVs values. Univariate normality was assessed using the Shapiro-Wilk test (p > 0.05), and multivariate normality was assessed using the Henze-Zirkler and Anderson-Darling tests (p < 0.05) to select the optimal test type (parametric or nonparametric). Nonparametric analysis was performed because the data were not normally distributed (p < 0.05). Statistical differences were assessed using Friedman’s test (p < 0.05), and multiple comparisons were performed using Nemenyi’s test (p < 0.05), which allows comparison of multiple methods while adjusting for type I error using the Tukey-Damico criterion. Additionally, a Cliff’s delta (δ) analysis was performed to assess the magnitude differences in performance metrics (Performance, Sensitivity, Specificity) measures between pairs of methods, complementing Nemenyi’s test, as a way to identify which pairs differ. We applied the aforementioned strategy independently within the frequency band and separately for the six-group and three-group representative time series, as well as for the multivariate extension of bivariate estimators.

At the second level, we applied an Aligned Rank Transform (ART) ANOVA, a nonparametric factorial method equivalent to 2-way ANOVA, to analyze data when normality or homoscedasticity assumptions were violated. The first factor in our analysis was the frequency bands, and the second factor was either the six methods for estimating the representative time series or the three methods for the multivariate extensions of bivariate connectivity estimators. With this approach, we aimed to identify any effects of frequency bands in our findings and to assess synergistic interactions between frequency bands and methods in both scenarios. We followed the same analysis in both datasets.

At the third level, we estimated the mean performance, sensitivity, and specificity for the first dataset and PDiVs for the second dataset of each of the 240 estimators across the six methods used to estimate the representative time series, or across the three methods used to estimate the multivariate extensions of bivariate connectivity estimators. This process led to a vector of 240 performance values across a) the six methods used to estimate the representative time series, and b) across the three methods used to estimate the multivariate extensions of bivariate connectivity estimators. The dimensionality of the multivariate estimators was maintained at seven. Statistical differences were assessed using the Wilcoxon rank sum test (p < 0.05, Bonferroni corrected).

Additionally, a Cliff’s delta (δ) analysis was performed to assess the magnitude differences in performance metrics for the first dataset (Performance, Sensitivity, Specificity) and in PDiVs for the second dataset between pairs of the three approaches, complementing Nemenyi’s test, as a way to identify which pairs differ.

Finally, we applied the Wilcoxon rank-sum test (p < 0.05) to the sparsity levels of FBNs between healthy controls and MCI subjects, independently for each connectivity estimator, across the three methodologies and frequency bands. We conducted this analysis to ensure that classification performance was not driven by differences in sparsity levels between the two groups. Similarly, we applied the Wilcoxon rank-sum test (p < 0.05) to the sparsity levels of FBNs between the three pairs of the three sessions.

### 3.4 Software tools

Pairwise interactions: A Python-based package called pyspi [74], which includes implementations of all 237 statistics on pairwise interactions (SPIs). This software allows users to quantify interactions between pairs of time series. Any interested reader can access the Python package at the following link:

https://pypi.org/project/pyspi/

The remaining three connectivity estimators (iPLV, ciPLV, CCc) can be downloaded from the following link:

https://github.com/stdimitr/fast_estimation_dynamic_functional_brain_networks.

Multivariate interactions: Multivariate connectivity estimators used here can be downloaded from the following link:

https://github.com/RikHenson/MultivarCon/.

Portrait divergence (PID): A Python implementation of PID for network comparisons can be downloaded from the following link:

https://github.com/bagrow/network-portrait-divergence.

Violin Plots: Implementation of Violin plots can be downloaded from the following link:

Violinplot-Matlab - File Exchange - MATLAB Central [75] Statistical analysis: All analyses were conducted in R v.4.5.2.

OMST: The implementation of Orthogonal Minimal Spanning Trees (OMSTs) in Python can be found here : https://github.com/makism/dyconnmap [76], and in MATLAB : https://github.com/stdimitr/topological_filtering_networks [66,67]

## 4. Results

### 4.1 The Impact of ROI extraction method and FC estimator on the Classification Performance between HC-MCI

#### 4.1.1 The effect of approaches based on representative time series and the multivariate extension of bivariate connectivity measures

Fig. 3 illustrates the classification performance of the kNN classifier using the PID metric on brain networks derived from healthy controls and MCI subjects. Results are presented across the frequency bands and the six approaches, based on representative time series, multivariate extensions of bivariate connectivity measures, and multivariate connectivity estimators. Below, we summarize the statistical tests used to evaluate classification performance. Findings on sensitivity and specificity, along with a similar statistical analysis, are presented in the supp. material.

**Fig. 3.**
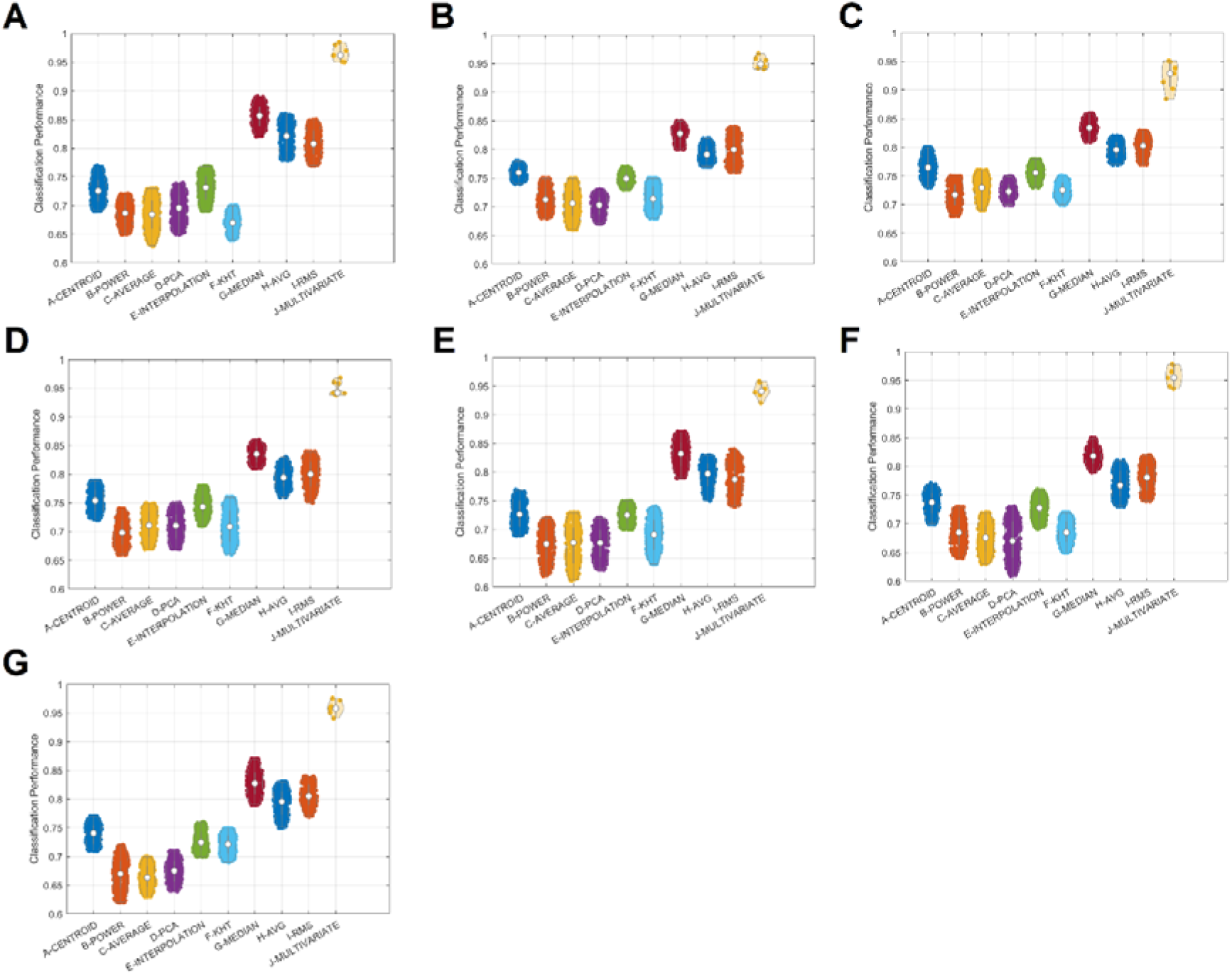
Classification performance of the machine learning approach applied over individual FBNs across the frequency bands, and the different approaches. (A-G : δ,θ,α_1_,α_2_,β_1_,β_2_,γ frequency bands)

Table 2 summarizes the results of Friedman’s test for the six approaches, based on representative time series from each frequency band. Results from Nemenyi’s test and Cliff’s delta are presented in supp. material for each frequency band. A consistent finding across frequency bands is that the CENTROID approach outperformed the remaining five methods, with small to moderate effect sizes (see Cliff’s δ values in the supp. material).

**Table 2.**
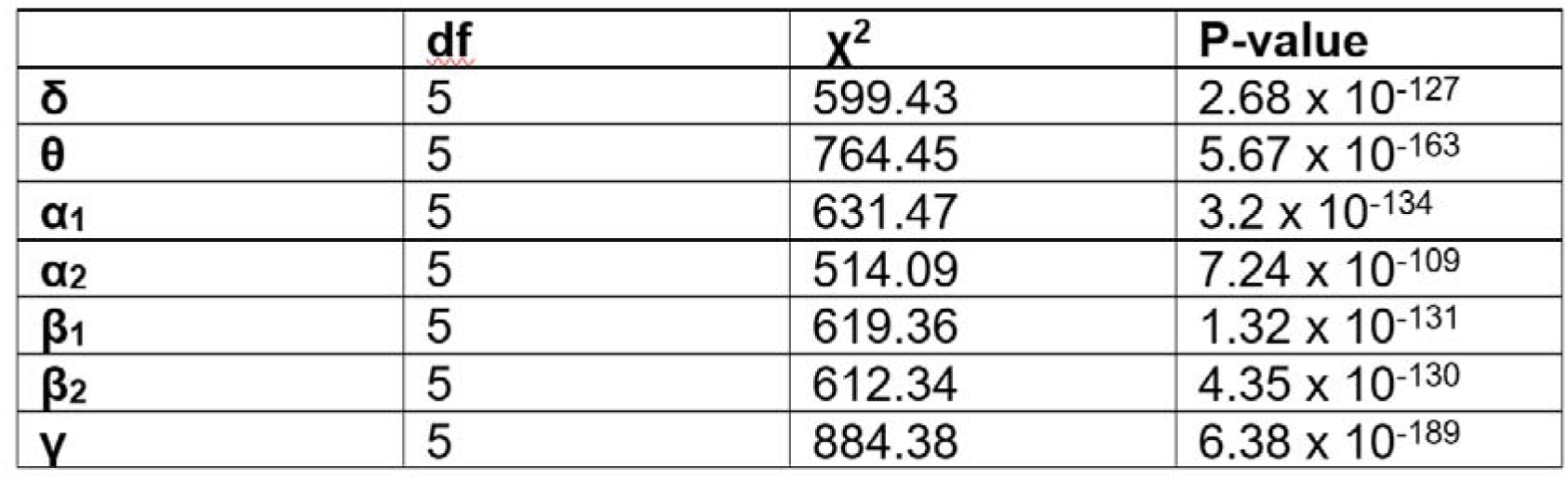
Summary of Friedman’s test per frequency band and across the six approaches for representative time series.

Table 3 summarizes the results of Friedman’s test for the three approaches and for the multivariate extensions of the bivariate connectivity estimators across frequency bands. Results from Nemenyi’s test and Cliff’s delta are presented in the supp. material for each frequency band. A consistent finding across frequency bands is that the MEDIAN approach outperformed the other two methods, with effect sizes ranging from small to large (see Cliff’s δ values in the supp. material).

**Table 3.** Summary of Friedman’s test per frequency band and across the three approaches for multivariate extensions of bivariate connectivity estimators.

|  | **df** | **$\chi^2$** | **P-value** |
| --- | --- | --- | --- |
| **$\delta$** | 2 | 222.3 | $5.34 \times 10^{-49}$ |
| **$\theta$** | 2 | 192.2 | $1.29 \times 10^{-42}$ |
| **$\alpha_1$** | 2 | 310.26 | $4.24 \times 10^{-68}$ |
| **$\alpha_2$** | 2 | 258.36 | $7.91 \times 10^{-57}$ |
| **$\beta_1$** | 2 | 178.61 | $1.64 \times 10^{-39}$ |
| **$\beta_2$** | 2 | 287.66 | $3.43 \times 10^{-63}$ |
| **$\gamma$** | 2 | 162.11 | $6.28 \times 10^{-36}$ |

We applied the Aligned Rank Transform (ART) ANOVA, a nonparametric factorial method equivalent to 2-way ANOVA, to assess the potential effects of frequency bands, the six approaches for estimating representative time series, the three approaches for the multivariate extension of bivariate estimators, and their interactions on classification performance. Our findings found only an effect of the methods and no effect of the frequency bands and their interactions (df = 5, F = 1222.15, p = 2.21 x 10^-16^, effect of the six approaches for estimating the representative time series, and df = 2, F = 912.43, p = 3.96 x 10^-14^, effect of the three approaches for the multivariate extension of the bivariate connectivity estimators).

#### 4.1.2 The performance of Multivariate estimators vs the approaches of representative time series and the multivariate extension of bivariate connectivity measures

Table 4 summarizes the statistical tests applied to the classification performance across pairs of methodologies and frequency bands. Our results show a significant improvement in classification performance between the multivariate estimator and the bivariate connectivity estimators compared with approaches for estimating representative time series, and further improvement when employing the multivariate estimators. We found similar results for the sensitivity and the specificity (see supp material).

**Table 4.** Summary of p-values derived from the application of the Wilcoxon rank sum test between pairs of methodologies and across the frequency bands for the classification performance. 1 refers to approaches for estimating representative time series, 2 to the multivariate extension of bivariate connectivity measures, and 3 to multivariate estimators.

|  | **1 vs 2** | **1 vs 3** | **2 vs 3** |
| --- | --- | --- | --- |
| $\delta$ | $5.42 \times 10^{-79}$ | $4.42 \times 10^{-7}$ | $5.44 \times 10^{-6}$ |
| $\theta$ | $3.72 \times 10^{-74}$ | $6.72 \times 10^{-7}$ | $6.23 \times 10^{-7}$ |
| $\alpha_1$ | $3.39 \times 10^{-77}$ | $5.38 \times 10^{-6}$ | $1.38 \times 10^{-6}$ |
| $\alpha_2$ | $5.64 \times 10^{-78}$ | $6.25 \times 10^{-6}$ | $5.83 \times 10^{-5}$ |
| $\beta_1$ | $4.29 \times 10^{-76}$ | $5.24 \times 10^{-5}$ | $2.36 \times 10^{-5}$ |
| $\beta_2$ | $3.05 \times 10^{-74}$ | $3.82 \times 10^{-5}$ | $5.71 \times 10^{-7}$ |
| $\gamma$ | $4.72 \times 10^{-75}$ | $5.27 \times 10^{-8}$ | $5.98 \times 10^{-6}$ |

#### 4.1.3 Statistical Analysis on the Sparsity Levels of Functional Brain Networks

We observed sparsity levels ranging from 11.11% (5 OMSTs - 5*89/4005) to 17.78% (8 OMSTs - 8*89/4005) across FC estimators, adopted methods, frequency bands, and subjects. Our statistical analysis of sparsity levels between the two groups of subjects didn’t reveal any statistically significant differences across FC estimators, methods, or frequency bands (Wilcoxon rank-sum test, p > 0.05).

### 4.2 The Impact of ROI extraction method and FC estimator on the Repeatability of Functional Brain Network (FBN) Topologies

#### 4.2.1 The effect of approaches based on representative time series and the multivariate extension of bivariate connectivity measures

Fig. 4 illustrates the repeatability of FBN topologies measured with PID estimated between pairs of sessions and averaged across subjects. Results are presented across the frequency bands and the six approaches, based on representative time series, multivariate extensions of bivariate connectivity measures, and multivariate connectivity estimators. Below, we summarize the statistical tests used to evaluate the repeatability.

**Fig. 4.**
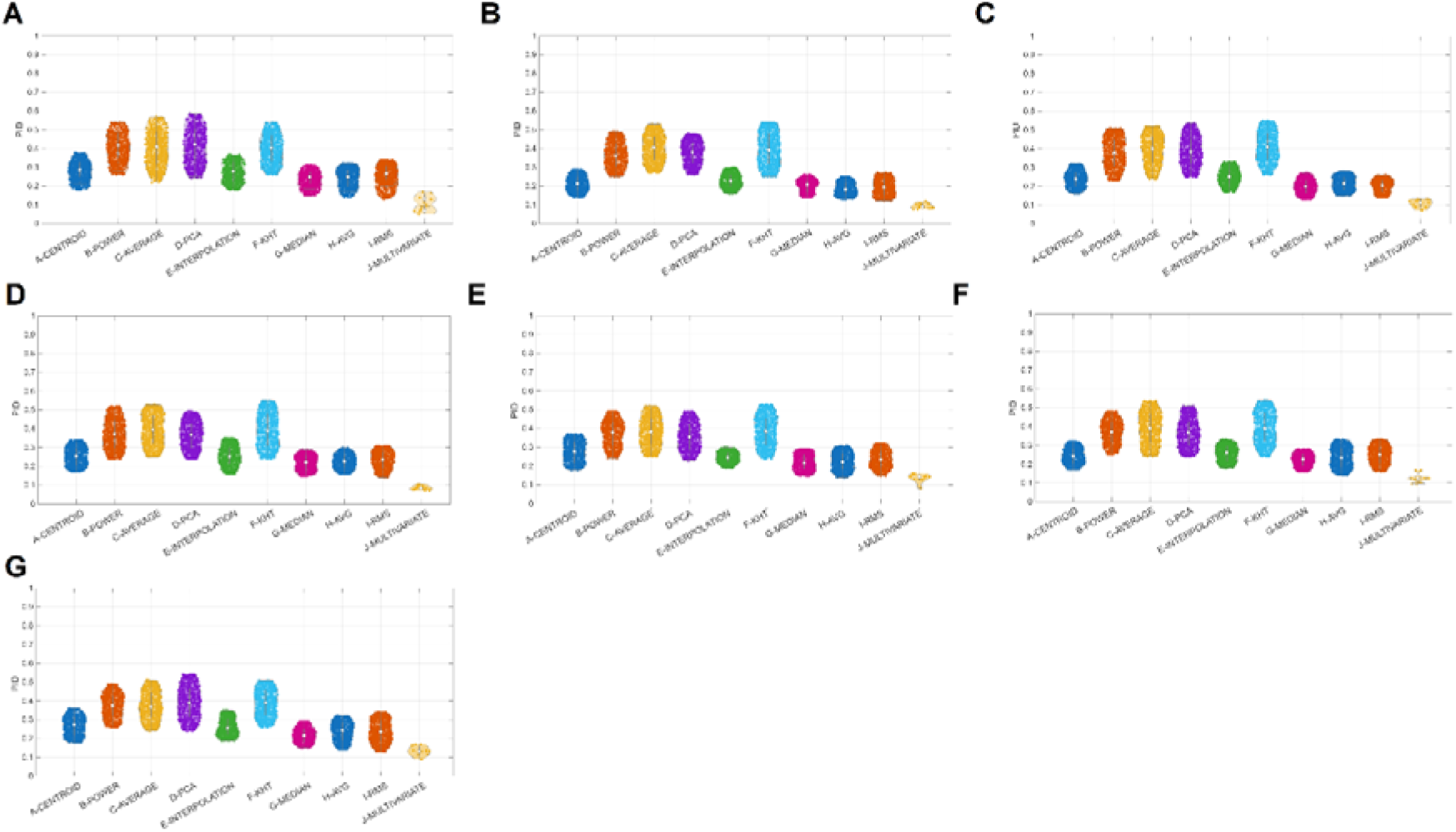
Repeatability of FBNs topologies between sessions estimated per individual and measured with PDiV across the frequency bands, and the different approaches. (A-G : δ,θ,α_1_,α_2_,β_1_,β_2_,γ frequency bands)

Table 5 summarizes the results of Friedman’s test for the six approaches, based on representative time series from each frequency band. Results from Nemenyi’s test and Cliff’s delta are presented in supp. material for each frequency band. A consistent finding across frequency bands is that the CENTROID approach outperformed the remaining five methods, with small to moderate effect sizes (see Cliff’s δ values in the supp. material).

**Table 5.** Summary of Friedman’s test per frequency band and across the six approaches for representative time series.

| | *df* | $\chi^2$ | *P-value* |
| --- | --- | --- | --- |
| $\delta$ | 5 | 482.3 | $5.26 \times 10^{-102}$ |
| $\theta$ | 5 | 772.27 | $1.44 \times 10^{-164}$ |
| $\alpha_1$ | 5 | 645.45 | $3.04 \times 10^{-137}$ |
| $\alpha_2$ | 5 | 486.82 | $5.58 \times 10^{-103}$ |
| $\beta_1$ | 5 | 517.67 | $1.22 \times 10^{-109}$ |
| $\beta_2$ | 5 | 594.66 | $2.87 \times 10^{-126}$ |
| $\gamma$ | 5 | 480.95 | $1.03 \times 10^{-101}$ |

Table 6 summarizes the results of Friedman’s test for the three approaches and for the multivariate extensions of the bivariate connectivity estimators across frequency bands. Results from Nemenyi’s test and Cliff’s delta are presented in the supp. material for each frequency band. A consistent finding across frequency bands is that the MEDIAN approach outperformed the other two methods, with effect sizes ranging from small to large (see Cliff’s δ values in the supp. material).

**Table 6.** Summary of Friedman’s test per frequency band and across the three approaches for multivariate extensions of bivariate connectivity estimators.

| | *df* | $\chi^2$ | *P-value* |
| --- | --- | --- | --- |
| $\delta$ | 2 | 29.78 | $3.42 \times 10^{-7}$ |
| $\theta$ | 2 | 13.41 | 0.0012 |
| $\alpha_1$ | 2 | 15.83 | 0.0004 |
| $\alpha_2$ | 2 | 4.41 | 0.11 |
| $\beta_1$ | 2 | 7.68 | 0.02 |
| $\beta_2$ | 2 | 35.76 | $1.71 \times 10^{-8}$ |
| $\gamma$ | 2 | 9.86 | 0.0072 |

We applied the Aligned Rank Transform (ART) ANOVA, a nonparametric factorial method equivalent to 2-way ANOVA, to assess the potential effects of frequency bands, the six approaches for estimating representative time series, the three approaches for the multivariate extension of bivariate estimators, and their interactions on repeatability. Our findings found only an effect of the methods and no effect of the frequency bands and their interactions (df = 5, F = 1466.57, p = 2.22 x 10^-308^, effect of the six approaches for estimating the representative time series, and df = 2, F = 241.22, p = 2.22 x 10^-308^, effect of the three approaches for the multivariate extension of the bivariate connectivity estimators). P-values reached MATLAB precision limits.

#### 4.2.2 The performance of Multivariate estimators vs the approaches of representative time series and the multivariate extension of bivariate connectivity measures

Table 7 summarizes the statistical tests applied to the repeatability across pairs of methodologies and frequency bands. Our results show a significant improvement in classification performance between the multivariate estimator and the bivariate connectivity estimators compared with approaches for estimating representative time series, and further improvement when employing the multivariate estimators. We found similar results for the sensitivity and the specificity (see supp material).

**Table 7.** Summary of p-values derived from the application of the Wilcoxon rank sum test between pairs of methodologies and across the frequency bands for the repeatability measurements. 1 refers to approaches for estimating representative time series, 2 to the multivariate extension of bivariate connectivity measures, and 3 to multivariate estimators.

|  | **1 vs 2** | **1 vs 3** | **2 vs 3** |
| --- | --- | --- | --- |
| $\delta$ | $4.12 \times 10^{-80}$ | $6.62 \times 10^{-6}$ | $6.62 \times 10^{-6}$ |
| $\theta$ | $3.64 \times 10^{-72}$ | $5.83 \times 10^{-7}$ | $6.67 \times 10^{-6}$ |
| $\alpha_1$ | $3.82 \times 10^{-71}$ | $5.05 \times 10^{-5}$ | $1.89 \times 10^{-7}$ |
| $\alpha_2$ | $4.34 \times 10^{-75}$ | $6.72 \times 10^{-7}$ | $5.24 \times 10^{-6}$ |
| $\beta_1$ | $5.37 \times 10^{-70}$ | $4.53 \times 10^{-6}$ | $3.77 \times 10^{-5}$ |
| $\beta_2$ | $3.67 \times 10^{-72}$ | $4.12 \times 10^{-5}$ | $4.92 \times 10^{-5}$ |
| $\gamma$ | $4.24 \times 10^{-71}$ | $5.62 \times 10^{-8}$ | $5.14 \times 10^{-6}$ |

#### 4.2.3 Statistical Analysis on the Sparsity Levels of Functional Brain Networks

We observed sparsity levels ranging from 11.11% (5 OMSTs - 5*89/4005) to 17.78% (8 OMSTs - 8*89/4005) across FC estimators, adopted methods, frequency bands, and subjects. Our statistical analysis of sparsity levels between the two groups of subjects didn’t reveal any statistically significant differences across FC estimators, methods, or frequency bands (Wilcoxon rank-sum test, p > 0.05).

## 5. Discussion

Extensive neuroimaging research utilizing magnetoencephalography and electroencephalography (M/EEG) seeks to identify reliable functional connectomic biomarkers of brain function and its disorders. However, this process is impeded by the many arbitrary methodological choices required [32,77].

The results of this study indicate that the selection of dimensionality reduction methods significantly influences the resulting connectivity measures. This effect was observed in both MCI-HC and test-retest datasets. Notably, the choice of extraction method produced substantial differences in the final FBN topology. The multivariate extensions of bivariate FC estimation (aggregation procedures : MEAN, MEDIAN, RMS) applied after computing connectivity produced more consistent results across both in vivo datasets compared to the dimensionality reduction approaches (CENTROID, POWER, AVERAGE, PCA, INTERPOLATION, KHT). The multivariate extensions of bivariate FC estimation approaches are computationally demanding but enable the integration of information from all elements within a given brain parcel, resulting in a more comprehensive estimation of FC and the derived FBN topology [24,32,38,39,45,77–79]. Multivariate FC estimators that simultaneously handle pairs of voxel time series between every pair of parcels further improved the estimation of FC and the overall FBN topology in both MEG resting-state datasets.

Functional connectivity (FC) is the primary framework for understanding how the brain functions at rest and during task performance, and how brain dysfunction occurs in various brain disorders and diseases. Brain recordings using EEG and MEG are acquired from outside the head, and estimating virtual source activity yields highly redundant brain activity. This redundancy is a consequence of recording brain activity with a few tens (EEG) or hundreds (MEG) sensors, and estimating virtual sources of a few thousand [16], and hinders accurate FC estimation. This problem is solved by adopting an atlas (e.g., AAL here [25]) that maps voxel-based time series to anatomically parcellated brain regions on the virtual source space [35]. However, estimating FC between pairs of brain regions is not trivial. A common approach for the construction of a brain network is: 1) the adoption of a method for the estimation of the representative time series per brain area from a set of voxel time series as it is defined by the adopted atlas [28–33], 2) the selection of an FC estimator, and 3) the quantification of functional strength between every pair of brain regions with the selected FC estimator [33]. This dimensionality-reduction approach requires selecting an appropriate method to estimate a representative time series, but it doesn’t capture the full information available between every pair of brain areas. Additionally, this approach overlooks the fact that brain regions differ markedly in size and shape. This heterogeneity suggests that using the same method to generate representative time series for each region to estimate FC between them is inappropriate.

To demonstrate the superiority of the representative free approaches, we estimated FC between brain regions using A) an exhaustive set of FC estimators paired with methods that define representative time series per brain area [28–33], widely used in the literature, and B) the multivariate extensions of bivariate FC estimators that use all voxel time series within the parcellated brain areas in a pair- wise fashion [34–36,38,49–52,80]. Additionally, we compared the two methods of estimating FC between a pair of brain regions to a set of multivariate connectivity estimators that simultaneously analyze brain activity across all voxel time series spatially constrained in the two brain areas [34–36,38,49,50,57,58,60,61,64]. These two alternative methods eliminate the need to estimate a representative time series for each brain region. Our framework was evaluated under two scenarios using MEG resting-state recordings : A) the classification performance between healthy controls and MCI subjects, and B) the repeatability of FBN topologies between test-retest sessions. Both MEG datasets are freely available.

The brain’s networked architecture facilitates synchrony among neuronal populations. These communication patterns are mapped using functional imaging such as EEG, MEG, and fMRI, resulting in functional connectivity (FC) networks. Although a small set of FC estimators is commonly employed particularly in MEG studies, a variety of pairwise interaction statistics are available in the scientific literature. We used a library of 240 pairwise statistics to capture FC in a pair-wise fashion across the 90 ROIs of the AAL tabulated in FBN of size 90 x 90 (2D tensor). We strongly believe that this study is a strong proof of principle showing the relevance of the aggregation procedures in studying resting state connectivity, similarly to rs-fMRI [44]. On the top, we underscore that our findings are independent of the choice of specific FC estimators supporting further their universal acceptance from the neuroscientists.

Our results demonstrated that the multivariate extension of bivariate FC estimators (representative-free approach), which summarizes pairwise FC strength across all voxels of two brain areas, and accurate multivariate estimators that consider pairs of region-wise voxel time series at once, clearly outperform bivariate FC estimators based on representative time series. Multivariate connectivity estimators outperformed both approaches (representative time series with bivariate FC estimators and multivariate extensions of bivariate FC estimators) in classifying healthy controls versus MCI using MEG resting-state recordings (Fig.3), and also on the repeatability of FBN topologies (Fig.4). Our findings were consistent across the frequency bands studied. A reasonable explanation of the improved performance of the two alternative approaches compared to the ROI extraction strategies is the better manipulation of the voxel-wise time-series across every pair of ROIs independently of their size.

Significant and widespread differences in connectivity outcomes were observed in both tasks across ROI dimensionality reduction approaches. Therefore, caution is necessary when comparing connectivity patterns across studies that use different ROI extraction strategies, since even minor methodological variations can result in substantial discrepancies (Fig.3 & Fig.4).

It is important to note that our analysis focused on A) classifying FBN topologies between healthy controls and MCI subjects, and B) on the repeatability of FBN between test-retest sessions using MEG resting-state recordings. We managed FBNs as 2D tensors of size [90 x 90] focusing on the whole FBN topology. Complementarily, we systematically evaluated the performance of an extreme set of FC estimators to independently support our findings, irrespective of the small subset commonly used in EEG/MEG FC analysis. Additionally, we didn’t intend to distinguish FC estimators based on amplitude, phase, or time/frequency domains in terms of their performance in both tasks. We aimed to underscore the significant effect of processing steps, such as the way we can extract and use ROI-based brain activity on the voxel-level, on the FC analysis and the resulting FBNs.

In representative time-series methods, the centroid consistently outperformed the other studied approaches across frequency bands, followed by interpolation and KHT. In the multivariate extension of bivariate FC estimators, MEDIAN demonstrated superior classification performance compared with AVERAGE and RMS, while it showed an inconsistent dependency on frequency bands. Multivariate FC estimators exhibited consistently high classification performance across both approaches and in both tasks. Our findings were not driven by any statistically significant difference in the sparsity levels of FBNs between the two groups (classification study), and between the sessions (test-retest study).

Here, we employed the AAL atlas, which comprises 90 brain areas and 2.459 source positions. This configuration yields 4.005 area-area pairs and a very large number of virtual-source-to-virtual-source pairs. The multivariate approaches require computing all source-to-source functional connectivity strengths, which dramatically increases computational demands. However, these computationally demanding tasks can be processed on a standard personal computer and, in large population studies, on a cluster.

### 5.1 Limitations

There are several limitations to the current study. First, the results may not be generalizable to all possible strategies, as this was beyond the study’s scope. The analysis focused on a set of common and practical scenarios. In fairness, some limitations of this work must be highlighted. First, the analysis involved specific processing steps for FBNs using the fixed atlas AAL highly used in MEG analysis. The effects of denoising algorithms and of selecting an alternative source-localization algorithm, using the same or an alternative atlas, could affect the FC analysis in either direction. We demonstrated in rs-fMRI that the synergy among data-processing steps is critical for constructing a reproducible and consistent functional connectome (Luppi et al., [42]). Here, we validated the superiority of two approaches over the trivial approach of estimating a representative time series for each brain area in a classification task between healthy controls and MCI, and in a test-retest study. Further exploratory studies are needed in this direction, using datasets tailored to specific brain disorders and age ranges [6,9,81–84].

Another limitation is that the study only focused on resting-state data. While many studies have shown that some patterns of connectivity remain consistent between rest and tasks, recent literature suggests that individual variations should be considered [85]. Therefore, the generalizability of the results to task-based data cannot be considered straightforward, but demands further analysis.

We believe that this study is a strong proof of principle showing the relevance of the aggregation procedures in studying resting state connectivity like in rs-fMRI [44] . We showed that these results are independent of the choice of connectivity measures and go beyond phase, amplitude, distance, information-theoretic, causal, and spectral based metrics. A recent study explored this set of FC estimators on rs-fMRI datasets showing the correspondence of the derived FBN to : structure– function coupling, other neurophysiological networks, individual fingerprinting and brain–behavior prediction [54]. Apart from the exhaustive set of FC estimators, considered here, there are also FC estimation methods that should be investigated in future studies.

## 6. Conclusions

Based on our inter-area FC analysis of two MEG resting-state cohorts, one of HC and MCI subjects, and a second one on test-retest sessions in a virtual source space, our study showed that multivariate extensions of bivariate FC estimators are natural alternatives to combining representative time series for each brain area with bivariate FC estimators. In terms of classification performance, and repeatability, multivariate connectivity estimators outperformed both approaches (representative time series with bivariate FC estimators and multivariate extensions of bivariate FC estimators). The results demonstrated that representative time series were highly dependent on the selected method. Furthermore, both their performance and repeatability were significantly lower than those of alternative approaches, as indicated by whole-brain connectivity analysis. Our findings strongly discourage the authors’ use of representative time series. However, further studies are needed to evaluate this finding in additional clinical cohorts, across a broader age range, and in cognitive tasks.

## Author contributions

Stavros I Dimitriadis: conceptualization, supervision, data curation, methodology, formal analysis, software, writing (original draft, reviewing and editing), and performed the whole analysis.

## Abbreviations

MEG: Magnetoencephalography
EEG: Electroencephalography
kNN: k Nearest Neighbor classifier
PID: Portrait Divergence
OMST: Orthogonal Minimal Spanning Trees
HC: Healthy Controls
MCI: Mild Cognitive Impairment
MNE: minimum-norm estimate (MNE)
LORETA: low-resolution electromagnetic tomography
sLORETA: standardized LORETA
AAL: Automated Anatomical Labeling
LCMV: Linearly Constrained Minimum Variance
FCBN: functional connectivity brain networks (FCBN)
wICA: wavelet ICA

