## Supplementary Material for "Systematic evaluation of an exhaustive set of connectivity estimators in bivariate and multivariate modes for an improved virtual source MEG connectivity analysis for the study of resting state recordings"

1. **Statistical analysis and post hoc tests for the Classification Performance**

**1.1 Representative time series**

**STable 1** summarizes the results of Friedman's test for the six approaches, based on representative time series across the frequency bands. Results from Nemenyi's test and Cliff’s delta for each frequency band are summarized in **STable 2.**

**STable 1.** Results of Friedman’s Test for the six approaches based on representative time series across the frequency bands.

|  | **df** | **χ^2^** | **P-value** |
| --- | --- | --- | --- |
| **δ** | 5 | 599.43 | 2.68 x 10^-127^ |
| **θ** | 5 | 764.45 | 5.67 x 10^-163^ |
| **α_1_** | 5 | 631.47 | 3.2 x 10^-134^ |
| **α_2_** | 5 | 514.09 | 7.24 x 10^-109^ |
| **β_1_** | 5 | 619.36 | 1.32 x 10^-131^ |
| **β_2_** | 5 | 612.34 | 4.35 x 10^-130^ |
| **γ** | 5 | 884.38 | 6.38 x 10^-189^ |

**STable 2. (A – G ).** Summary of Nemenyi's test and Cliff’s delta for each frequency band.

A. **delta frequency**

Method 1	Method 2	LowerLimit	Difference	UpperLimit	Pvalue	Cliffs d

1	2	1.7217	2.2083	2.695	0	0

1	3	1.8175	2.3042	2.7908	0	0.13333

1	4	1.3008	1.7875	2.2742	0	-0.066667

1	5	-0.57418	-0.0875	0.39918	0.99572	0.2

1	6	2.6758	3.1625	3.6492	0	0.066667

2	3	-0.39085	0.095833	0.58251	0.99345	0.066667

2	4	-0.90751	-0.42083	0.065847	0.13486	-0.33333

2	5	-2.7825	-2.2958	-1.8092	0	0.13333

2	6	0.46749	0.95417	1.4408	3.4245e-07	-0.066667

3	4	-1.0033	-0.51667	-0.029987	0.029904	-0.4

3	5	-2.8783	-2.3917	-1.905	0	0.066667

3	6	0.37165	0.85833	1.345	7.4266e-06	-0.26667

4	5	-2.3617	-1.875	-1.3883	0	0.26667

4	6	0.88832	1.375	1.8617	6.0977e-15	0.26667

5	6	2.7633	3.25	3.7367	0	-0.2

B. **Theta frequency**

Method 1	Method 2	LowerLimit	Difference	UpperLimit	Pvalue	Cliffs d

1	2	1.7217	2.2083	2.695	0	-0.33333

1	3	1.8175	2.3042	2.7908	0	-0.33333

1	4	1.3008	1.7875	2.2742	0	-0.13333

1	5	-0.57418	-0.0875	0.39918	0.99572	-0.26667

1	6	2.6758	3.1625	3.6492	0	0.13333

2	3	-0.39085	0.095833	0.58251	0.99345	-0.066667

2	4	-0.90751	-0.42083	0.065847	0.13486	0.066667

2	5	-2.7825	-2.2958	-1.8092	0	0.066667

2	6	0.46749	0.95417	1.4408	3.4245e-07	0.33333

3	4	-1.0033	-0.51667	-0.029987	0.029904	0.26667

3	5	-2.8783	-2.3917	-1.905	0	0

3	6	0.37165	0.85833	1.345	7.4266e-06	0.53333

4	5	-2.3617	-1.875	-1.3883	0	-0.2

4	6	0.88832	1.375	1.8617	6.0977e-15	0.2

5	6	2.7633	3.25	3.7367	0	0.4

C. **Alpha1 frequency**

Method 1	Method 2	LowerLimit	Difference	UpperLimit	Pvalue	Cliffs d

1	2	1.7217	2.2083	2.695	0	0.2

1	3	1.8175	2.3042	2.7908	0	0.2

1	4	1.3008	1.7875	2.2742	0	0.86667

1	5	-0.57418	-0.0875	0.39918	0.99572	0.33333

1	6	2.6758	3.1625	3.6492	0	0.2

2	3	-0.39085	0.095833	0.58251	0.99345	0.13333

2	4	-0.90751	-0.42083	0.065847	0.13486	0.73333

2	5	-2.7825	-2.2958	-1.8092	0	-0.066667

2	6	0.46749	0.95417	1.4408	3.4245e-07	0.066667

3	4	-1.0033	-0.51667	-0.029987	0.029904	0.53333

3	5	-2.8783	-2.3917	-1.905	0	-0.2

3	6	0.37165	0.85833	1.345	7.4266e-06	0

4	5	-2.3617	-1.875	-1.3883	0	-0.66667

4	6	0.88832	1.375	1.8617	6.0977e-15	-0.53333

5	6	2.7633	3.25	3.7367	0	0.2

D. **Alpha2 frequency**

Method 1	Method 2	LowerLimit	Difference	UpperLimit	Pvalue	Cliffs d

1	2	1.7217	2.2083	2.695	0	-0.2

1	3	1.8175	2.3042	2.7908	0	-0.13333

1	4	1.3008	1.7875	2.2742	0	0.066667

1	5	-0.57418	-0.0875	0.39918	0.99572	-0.26667

1	6	2.6758	3.1625	3.6492	0	0.13333

2	3	-0.39085	0.095833	0.58251	0.99345	0

2	4	-0.90751	-0.42083	0.065847	0.13486	0.26667

2	5	-2.7825	-2.2958	-1.8092	0	0.066667

2	6	0.46749	0.95417	1.4408	3.4245e-07	0.53333

3	4	-1.0033	-0.51667	-0.029987	0.029904	0.46667

3	5	-2.8783	-2.3917	-1.905	0	0

3	6	0.37165	0.85833	1.345	7.4266e-06	0.26667

4	5	-2.3617	-1.875	-1.3883	0	-0.26667

4	6	0.88832	1.375	1.8617	6.0977e-15	0.066667

5	6	2.7633	3.25	3.7367	0	0.53333

E. **Beta 1 frequency**

Method 1	Method 2	LowerLimit	Difference	UpperLimit	Pvalue	Cliffs d

1	2	1.7217	2.2083	2.695	0	-0.2

1	3	1.8175	2.3042	2.7908	0	-0.33333

1	4	1.3008	1.7875	2.2742	0	0.066667

1	5	-0.57418	-0.0875	0.39918	0.99572	-0.2

1	6	2.6758	3.1625	3.6492	0	-0.2

2	3	-0.39085	0.095833	0.58251	0.99345	-0.13333

2	4	-0.90751	-0.42083	0.065847	0.13486	0.26667

2	5	-2.7825	-2.2958	-1.8092	0	0.2

2	6	0.46749	0.95417	1.4408	3.4245e-07	0.066667

3	4	-1.0033	-0.51667	-0.029987	0.029904	0.46667

3	5	-2.8783	-2.3917	-1.905	0	0.2

3	6	0.37165	0.85833	1.345	7.4266e-06	0.26667

4	5	-2.3617	-1.875	-1.3883	0	-0.2

4	6	0.88832	1.375	1.8617	6.0977e-15	0.066667

5	6	2.7633	3.25	3.7367	0	0.066667

F. **Beta 2 frequency**

Method 1	Method 2	LowerLimit	Difference	UpperLimit	Pvalue	Cliffs d

1	2	1.7217	2.2083	2.695	0	0.4

1	3	1.8175	2.3042	2.7908	0	0.2

1	4	1.3008	1.7875	2.2742	0	0.26667

1	5	-0.57418	-0.0875	0.39918	0.99572	0.4

1	6	2.6758	3.1625	3.6492	0	0.66667

2	3	-0.39085	0.095833	0.58251	0.99345	-0.2

2	4	-0.90751	-0.42083	0.065847	0.13486	-0.2

2	5	-2.7825	-2.2958	-1.8092	0	-0.2

2	6	0.46749	0.95417	1.4408	3.4245e-07	0.26667

3	4	-1.0033	-0.51667	-0.029987	0.029904	0.066667

3	5	-2.8783	-2.3917	-1.905	0	-0.066667

3	6	0.37165	0.85833	1.345	7.4266e-06	0.33333

4	5	-2.3617	-1.875	-1.3883	0	0

4	6	0.88832	1.375	1.8617	6.0977e-15	0.33333

5	6	2.7633	3.25	3.7367	0	0.46667

G. **Gamma frequency**

Method 1	Method 2	LowerLimit	Difference	UpperLimit	Pvalue	Cliff’s δ

1	2	1.7217	2.2083	2.695	0	-0.2

1	3	1.8175	2.3042	2.7908	0	-0.13333

1	4	1.3008	1.7875	2.2742	0	-0.13333

1	5	-0.57418	-0.0875	0.39918	0.99572	0

1	6	2.6758	3.1625	3.6492	0	0

2	3	-0.39085	0.095833	0.58251	0.99345	-0.066667

2	4	-0.90751	-0.42083	0.065847	0.13486	-0.066667

2	5	-2.7825	-2.2958	-1.8092	0	0.066667

2	6	0.46749	0.95417	1.4408	3.4245e-07	0.066667

3	4	-1.0033	-0.51667	-0.029987	0.029904	0

3	5	-2.8783	-2.3917	-1.905	0	0.13333

3	6	0.37165	0.85833	1.345	7.4266e-06	0.13333

4	5	-2.3617	-1.875	-1.3883	0	-0.066667

4	6	0.88832	1.375	1.8617	6.0977e-15	-0.066667

5	6	2.7633	3.25	3.7367	0	0

1.2 **Multivariate Approaches for bivariate measures**

**STable 3.** summarizes the results of Friedman's test for the three approaches of the multivariate extension of bivariate FC estimators across the frequency bands. Results from Nemenyi's test and Cliff’s delta for each frequency band are summarized in **STable 4.**

**STable 3.** Results of Friedman’s Test for the three approaches of the multivariate extension of bivariate FC estimators across the frequency bands.

|  | **df** | **χ^2^** | **P-value** |
| --- | --- | --- | --- |
| **δ** | 2 | 222.3 | 5.34 x 10^-49^ |
| **θ** | 2 | 192.2 | 1.29 x 10^-42^ |
| **α_1_** | 2 | 310.26 | 4.24 x 10^-68^ |
| **α_2_** | 2 | 258.36 | 7.91 x 10^-57^ |
| **β_1_** | 2 | 178.61 | 1.64 x 10^-39^ |
| **β_2_** | 2 | 287.66 | 3.43 x 10^-63^ |
| **γ** | 2 | 162.11 | 6.28 x 10^-36^ |

**STable 4. (A – G ).** Summary of Nemenyi's test and Cliff’s delta for each frequency band.

**A. Delta – frequency**

Method 1	Method 2	LowerLimit	Difference	UpperLimit	Pvalue Cliff’s δ

1	2	0.83605	1.05	1.2639	0	0.33

1	3	1.0611	1.275	1.4889	0	0

2	3	0.01105	0.225	0.43895	0.036528	-0.33

**B. Theta – frequency**

Method 1	Method 2	LowerLimit	Difference	UpperLimit	Pvalue Cliff’s δ

1	2	0.96105	1.175	1.3889	0	-0.667

1	3	0.78605	1	1.2139	0	-0.667

2	3	-0.38895	-0.175	0.03895	0.13384	0.33

**C. Alpha1 – frequency**

Method 1	Method 2	LowerLimit	Difference	UpperLimit	Pvalue Cliff’s δ

1	2	1.3027	1.5167	1.7306	0	-0.33

1	3	1.0069	1.2208	1.4348	0	0.33

2	3	-0.50978	-0.29583	-0.081884	0.0034169	0.33

D. **Alpha 2 – frequency**

Method 1	Method 2	LowerLimit	Difference	UpperLimit	Pvalue Cliff’s δ

1	2	1.1527	1.3667	1.5806	0	0

1	3	0.93188	1.1458	1.3598	0	0

2	3	-0.43478	-0.22083	-0.0068837	0.041192	-0.33

**E. Beta 1 – frequency**

Method 1	Method 2	LowerLimit	Difference	UpperLimit	Pvalue Cliff’s δ

1	2	0.85688	1.0708	1.2848	0	0

1	3	0.82772	1.0417	1.2556	0	-0.66

2	3	-0.24312	-0.029167	0.18478	0.94528	-0.33

**F. Beta 2 – frequency**

Method 1	Method 2	LowerLimit	Difference	UpperLimit	Pvalue Cliff’s δ

1	2	1.2569	1.4708	1.6848	0	-0.33

1	3	0.94022	1.1542	1.3681	0	-0.33

2 3	-0.53062	-0.31667	-0.10272	0.0015159	-0.33

**G.Gamma frequency**

Method 1	Method 2	LowerLimit	Difference	UpperLimit	Pvalue	Cliff’s δ

1	2	0.94022	1.1542	1.3681	0	0.33

1	3	0.48188	0.69583	0.90978	6.1863e-14	0

2	3	-0.67228	-0.45833	-0.24438	1.5391e-06	-0.33

1.3 **The classification performance of Multivariate estimators vs the approaches of representative time series and the multivariate extension of bivariate connectivity measures**

**STable 5** summarizes p-values derived from the application of the Wilcoxon rank sum test between pairs of methodologies and across the frequency bands for the classification performance

**STable 5.** Summary of p-values derived from the application of the Wilcoxon rank sum test between pairs of methodologies and across the frequency bands for the classification performance. 1 refers to approaches for estimating representative time series, 2 to the multivariate extension of bivariate connectivity measures, and 3 to multivariate estimators.

|  | **1 vs 2** | **1 vs 3** | **2 vs 3** |
| --- | --- | --- | --- |
| **δ** | 4.12 x 10^-75^ | 6.62 x 10^-6^ | 5.67 x 10^-5^ |
| **θ** | 1.34 x 10^-72^ | 5.41 x 10^-7^ | 6.11 x 10^-7^ |
| **α_1_** | 5.44 x 10^-73^ | 4.46 x 10^-6^ | 4.82 x 10^-6^ |
| **α_2_** | 4.39 x 10^-78^ | 3.37 x 10^-5^ | 5.72 x 10^-8^ |
| **β_1_** | 3.87 x 10^-69^ | 2.38 x 10^-5^ | 4.81 x 10^-7^ |
| **β_2_** | 2.48 x 10^-70^ | 4.52 x 10^-7^ | 5.93 x 10^-6^ |
| **γ** | 3.77 x 10^-81^ | 6.49 x 10^-8^ | 5.61 x 10^-5^ |

2. **Statistical analysis and post hoc tests for the Sensitivity**

**SFig.1** illustrates the sensitivity metric of the kNN classifier using the PID metric applied to brain networks derived from healthy controls and MCI. Results are presented across the frequency bands and the six approaches, based on representative time series, multivariate extensions of bivariate connectivity measures, and multivariate connectivity estimators.

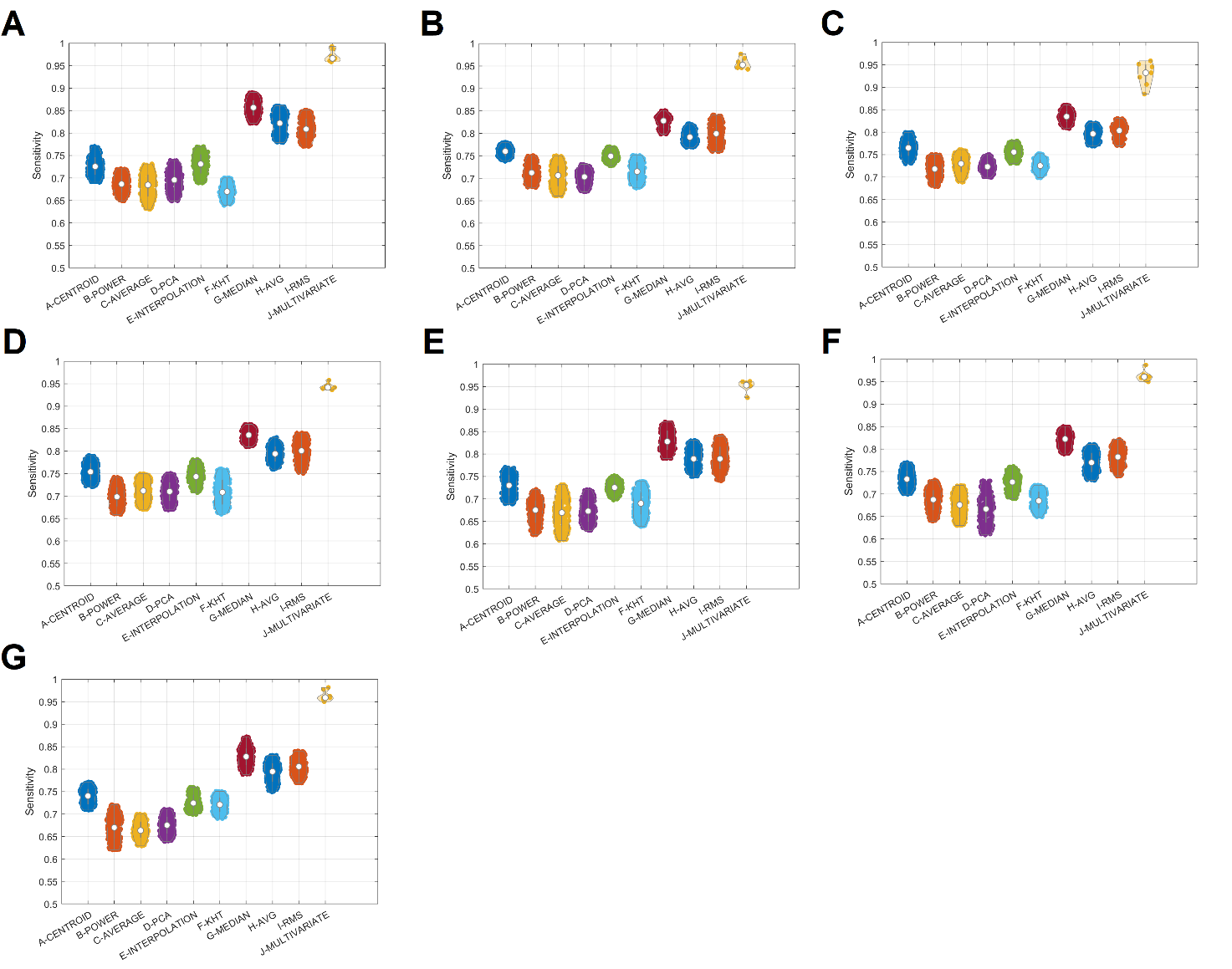

**SFig.1.** Sensitivity of the machine learning approach applied over individual functional brain networks across the frequency bands, and the different approaches. (A-G : δ,θ,α_1_,α_2_,β_1_,β_2_,γ frequency bands)

**2.1 Representative time series**

**STable 6** summarizes the results of Friedman's test for the six approaches, based on representative time series across the frequency bands. Results from Nemenyi's test and Cliff’s delta for each frequency band are summarized in **STable 7.**

**STable 6.** Results of Friedman’s Test for the six approaches based on representative time series across the frequency bands.

|  | **df** | **χ^2^** | **P-value** |
| --- | --- | --- | --- |
| **δ** | 5 | 598.67 | 3.91 x 10^-127^ |
| **θ** | 5 | 756.18 | 3.48 x 10^-161^ |
| **α_1_** | 5 | 629.67 | 7.86 x 10^-134^ |
| **α_2_** | 5 | 508.06 | 1.45 x 10^-107^ |
| **β_1_** | 5 | 616.24 | 6.26 x 10^-131^ |
| **β_2_** | 5 | 605.47 | 1.33 x 10^-128^ |
| **γ** | 5 | 879.67 | 6.66 x 10^-188^ |

**STable 7. (A – G ).** Summary of Nemenyi's test and Cliff’s delta for each frequency band.

**A.delta frequency**

Method 1	Method 2	LowerLimit	Difference	UpperLimit	Pvalue	Cliffs d

1	2	1.7383	2.225	2.7117	0	0.8037

1	3	1.8175	2.3042	2.7908	0	0.75666

1	4	1.3133	1.8	2.2867	0	0.61862

1	5	-0.58251	-0.095833	0.39085	0.99345	-0.081764

1	6	2.655	3.1417	3.6283	0	0.97252

2	3	-0.40751	0.079167	0.56585	0.99734	0.066771

2	4	-0.91168	-0.425	0.06168	0.12738	-0.1765

2	5	-2.8075	-2.3208	-1.8342	0	-0.82842

2	6	0.42999	0.91667	1.4033	1.1861e-06	0.42427

3	4	-0.99085	-0.50417	-0.017487	0.037221	-0.22782

3	5	-2.8867	-2.4	-1.9133	0	-0.78682

3	6	0.35082	0.8375	1.3242	1.3895e-05	0.25568

4	5	-2.3825	-1.8958	-1.4092	0	-0.66405

4	6	0.85499	1.3417	1.8283	3.4027e-14	0.51105

5 6	2.7508	3.2375	3.7242	0	0.97071

**B. Theta frequency**

Method 1	Method 2	LowerLimit	Difference	UpperLimit	Pvalue	Cliffs d

1	2	2.3925	2.8792	3.3658	0	0.96503

1	3	2.7633	3.25	3.7367	0	0.97326

1	4	3.0258	3.5125	3.9992	0	1

1	5	0.10499	0.59167	1.0783	0.007035	0.44038

1	6	2.28	2.7667	3.2533	0	0.95129

2	3	-0.11585	0.37083	0.85751	0.25119	0.19554

2	4	0.14665	0.63333	1.12	0.0028617	0.32109

2	5	-2.7742	-2.2875	-1.8008	0	-0.86328

2	6	-0.59918	-0.1125	0.37418	0.9863	-0.049651

3	4	-0.22418	0.2625	0.74918	0.64018	0.075732

3	5	-3.145	-2.6583	-2.1717	0	-0.88881

3	6	-0.97001	-0.48333	0.0033465	0.052827	-0.23787

4	5	-3.4075	-2.9208	-2.4342	0	-0.99895

4	6	-1.2325	-0.74583	-0.25915	0.00018268	-0.36266

5	6	1.6883	2.175	2.6617	0	0.82294

C. **Alpha1 frequency**

**Method 1	Method 2	LowerLimit	Difference	UpperLimit	Pvalue	Cliffs d**

1	2	2.6925	3.1792	3.6658	0	0.91224

1	3	1.9092	2.3958	2.8825	0	0.79773

1	4	2.3508	2.8375	3.3242	0	0.92106

1	5	-0.082513	0.40417	0.89085	0.16809	0.29205

1	6	2.1717	2.6583	3.145	0	0.8948

2	3	-1.27	-0.78333	-0.29665	6.5985e-05	-0.29501

2	4	-0.82835	-0.34167	0.14501	0.34184	-0.17109

2	5	-3.2617	-2.775	-2.2883	0	-0.86402

2	6	-1.0075	-0.52083	-0.034153	0.027759	-0.2416

3	4	-0.045013	0.44167	0.92835	0.10059	0.15722

3	5	-2.4783	-1.9917	-1.505	0	-0.70087

3	6	-0.22418	0.2625	0.74918	0.64018	0.087866

4	5	-2.92	-2.4333	-1.9467	0	-0.87469

4	6	-0.66585	-0.17917	0.30751	0.90118	-0.09463

5	6	1.7675	2.2542	2.7408	0	0.8356

D. **Alpha2 frequency**

**Method 1	Method 2	LowerLimit	Difference	UpperLimit	Pvalue	Cliffs d**

1	2	2.5217	3.0083	3.495	0	0.93114

1	3	1.8258	2.3125	2.7992	0	0.80757

1	4	1.905	2.3917	2.8783	0	0.83536

1	5	-0.03668	0.45	0.93668	0.088978	0.24568

1	6	1.8508	2.3375	2.8242	0	0.75471

2	3	-1.1825	-0.69583	-0.20915	0.00065691	-0.29693

2	4	-1.1033	-0.61667	-0.12999	0.0041337	-0.26796

2	5	-3.045	-2.5583	-2.0717	0	-0.86084

2	6	-1.1575	-0.67083	-0.18415	0.0012041	-0.21614

3	4	-0.40751	0.079167	0.56585	0.99734	0.030649

3	5	-2.3492	-1.8625	-1.3758	0	-0.69093

3	6	-0.46168	0.025	0.51168	0.99999	0.029324

4	5	-2.4283	-1.9417	-1.455	0	-0.72503

4	6	-0.54085	-0.054167	0.43251	0.99957	0.0061018

5	6	1.4008	1.8875	2.3742	0	0.63713

E. **Beta 1 frequency**

**Method 1	Method 2	LowerLimit	Difference	UpperLimit	Pvalue	Cliffs d**

1	2	2.2508	2.7375	3.2242	0	0.87657

1	3	2.2383	2.725	3.2117	0	0.82256

1	4	2.2508	2.7375	3.2242	0	0.86956

1	5	-0.48251	0.0041667	0.49085	1	0.10833

1	6	1.3592	1.8458	2.3325	0	0.66771

2	3	-0.49918	-0.0125	0.47418	1	0.049861

2	4	-0.48668	0	0.48668	1	-0.035321

2	5	-3.22	-2.7333	-2.2467	0	-0.93717

2	6	-1.3783	-0.89167	-0.40499	2.6416e-06	-0.3401

3	4	-0.47418	0.0125	0.49918	1	-0.076743

3	5	-3.2075	-2.7208	-2.2342	0	-0.85485

3	6	-1.3658	-0.87917	-0.39249	3.91e-06	-0.34045

4	5	-3.22	-2.7333	-2.2467	0	-0.92622

4	6	-1.3783	-0.89167	-0.40499	2.6416e-06	-0.32594

5	6	1.355	1.8417	2.3283	0	0.69079

F. **Beta 2 frequency**

**Method 1	Method 2	LowerLimit	Difference	UpperLimit	Pvalue	Cliffs d**

1	2	1.9633	2.45	2.9367	0	0.85275

1	3	2.3425	2.8292	3.3158	0	0.94121

1	4	2.6883	3.175	3.6617	0	0.88577

1	5	-0.03668	0.45	0.93668	0.088978	0.23584

1	6	1.9842	2.4708	2.9575	0	0.9113

2	3	-0.10751	0.37917	0.86585	0.22831	0.24222

2	4	0.23832	0.725	1.2117	0.00031478	0.31569

2	5	-2.4867	-2	-1.5133	0	-0.74344

2	6	-0.46585	0.020833	0.50751	1	0.013285

3	4	-0.14085	0.34583	0.83251	0.32794	0.13156

3	5	-2.8658	-2.3792	-1.8925	0	-0.86067

3	6	-0.84501	-0.35833	0.12835	0.28809	-0.24881

4	5	-3.2117	-2.725	-2.2383	0	-0.82793

4	6	-1.1908	-0.70417	-0.21749	0.00053405	-0.31778

5	6	1.5342	2.0208	2.5075	0	0.79937

G. **Gamma frequency**

**Method 1	Method 2	LowerLimit	Difference	UpperLimit	Pvalue	Cliffs d**

1	2	2.8758	3.3625	3.8492	0	0.98135

1	3	3.1633	3.65	4.1367	0	1

1	4	2.6092	3.0958	3.5825	0	0.99934

1	5	0.12999	0.61667	1.1033	0.0041337	0.36513

1	6	0.43832	0.925	1.4117	9.0382e-07	0.51095

2	3	-0.19918	0.2875	0.77418	0.54279	0.083647

2	4	-0.75335	-0.26667	0.22001	0.62413	-0.12559

2	5	-3.2325	-2.7458	-2.2592	0	-0.92723

2	6	-2.9242	-2.4375	-1.9508	0	-0.87312

3	4	-1.0408	-0.55417	-0.067487	0.01492	-0.27497

3	5	-3.52	-3.0333	-2.5467	0	-0.99983

3	6	-3.2117	-2.725	-2.2383	0	-0.98044

4	5	-2.9658	-2.4792	-1.9925	0	-0.96095

4	6	-2.6575	-2.1708	-1.6842	0	-0.89756

5	6	-0.17835	0.30833	0.79501	0.46215	0.19243

2.2 **Multivariate Approaches for bivariate measures**

**STable 8.** summarizes the results of Friedman's test for the three approaches of the multivariate extension of bivariate FC estimators across the frequency bands. Results from Nemenyi's test and Cliff’s delta for each frequency band are summarized in **STable 9.**

**STable 8.** Results of Friedman’s Test for the three approaches of the multivariate extension of bivariate FC estimators across the frequency bands.

|  | **df** | **χ^2^** | **P-value** |
| --- | --- | --- | --- |
| **δ** | 2 | 220.68 | 1.20 x 10^-48^ |
| **θ** | 2 | 194.91 | 4.74 x 10^-43^ |
| **α_1_** | 2 | 304.23 | 8.67 x 10^-67^ |
| **α_2_** | 2 | 261.56 | 1.59 x 10^-57^ |
| **β_1_** | 2 | 172.43 | 3.60 x 10^-38^ |
| **β_2_** | 2 | 288.96 | 1.79 x 10^-63^ |
| **γ** | 2 | 162.3 | 5.71 x 10^-36^ |

**STable 9. (A – G ).** Summary of Nemenyi's test and Cliff’s delta for each frequency band.

**A. Delta – frequency**

Method 1	Method 2	LowerLimit	Difference	UpperLimit	Pvalue	Cliffs d

1	2	0.82355	1.0375	1.2514	0	0.71618

1	3	1.0611	1.275	1.4889	0	0.86067

2	3	0.02355	0.2375	0.45145	0.025146	0.24432

**B. Theta – frequency**

Method 1	Method 2	LowerLimit	Difference	UpperLimit	Pvalue	Cliffs d

1	2	0.96522	1.1792	1.3931	0	0.86084

1	3	0.79438	1.0083	1.2223	0	0.59055

2	3	-0.38478	-0.17083	0.043116	0.14703	-0.11942

**C. Alpha1 – frequency**

Method 1	Method 2	LowerLimit	Difference	UpperLimit	Pvalue	Cliffs

1	2	1.2861	1.5	1.7139	0	0.95268

1	3	0.99855	1.2125	1.4264	0	0.86252

2	3	-0.50145	-0.2875	-0.07355	0.0046609	-0.22005

D. **Alpha 2 – frequency**

Method 1	Method 2	LowerLimit	Difference	UpperLimit	Pvalue	Cliffs d

1	2	1.1652	1.3792	1.5931	0	0.92085

1	3	0.93188	1.1458	1.3598	0	0.75704

2	3	-0.44728	-0.23333	-0.019384	0.02854	-0.13602

**E. Beta 1 – frequency**

Method 1	Method 2	LowerLimit	Difference	UpperLimit	Pvalue	Cliffs d

1	2	0.84438	1.0583	1.2723	0	0.72779

1	3	0.80272	1.0167	1.2306	0	0.66423

2	3	-0.25562	-0.041667	0.17228	0.89154	-0.020188

**F. Beta 2 – frequency**

Method 1	Method 2	LowerLimit	Difference	UpperLimit	Pvalue	Cliffs d

1	2	1.2652	1.4792	1.6931	0	0.91771

1	3	0.93188	1.1458	1.3598	0	0.80656

2	3	-0.54728	-0.33333	-0.11938	0.00076211	-0.26893

**G.Gamma frequency**

Method 1	Method 2	LowerLimit	Difference	UpperLimit	Pvalue	Cliffs d

1	2	0.93605	1.15	1.3639	0	0.70798

1	3	0.51105	0.725	0.93895	4.399e-15	0.51967

2	3	-0.63895	-0.425	-0.21105	9.6372e-06	-0.32183

We applied the Aligned Rank Transform (ART) ANOVA, a nonparametric factorial method equivalent to 2-way ANOVA, to assess the potential effects of frequency bands, the six approaches for estimating representative time series, the three approaches for the multivariate extension of bivariate estimators, and their interactions on sensitivity. Our findings found only an effect of the methods and no effect of the frequency bands and their interactions (df = 5, F = 1128.42, p = 3.42 x 10^-15^, effect of the six approaches for estimating the representative time series, and df = 2, F = 878.62, p = 4.72 x 10^-13^, effect of the three approaches for the multivariate extension of the bivariate connectivity estimators).

2.3 **The classification performance of Multivariate estimators vs the approaches of representative time series and the multivariate extension of bivariate connectivity measures**

**STable 10** summarizes p-values derived from the application of the Wilcoxon rank sum test between pairs of methodologies and across the frequency bands for the sensitivity measurements.

**STable 10.** Summary of p-values derived from the application of the Wilcoxon rank sum test between pairs of methodologies and across the frequency bands for the sensitivity measurements. 1 refers to approaches for estimating representative time series, 2 to the multivariate extension of bivariate connectivity measures, and 3 to multivariate estimators.

|  | **1 vs 2** | **1 vs 3** | **2 vs 3** |
| --- | --- | --- | --- |
| **δ** | 4.12 x 10^-78^ | 6.62 x 10^-7^ | 6.27 x 10^-8^ |
| **θ** | 3.56 x 10^-74^ | 5.43 x 10^-8^ | 7.59 x 10^-7^ |
| **α_1_** | 2.34 x 10^-75^ | 7.12 x 10^-9^ | 4.29 x 10^-6^ |
| **α_2_** | 4.83 x 10^-76^ | 8.36 x 10^-6^ | 7.27 x 10^-5^ |
| **β_1_** | 3.95 x 10^-77^ | 4.58 x 10^-5^ | 5.28 x 10^-6^ |
| **β_2_** | 2.74 x 10^-72^ | 3.74 x 10^-7^ | 4.34 x 10^-8^ |
| **γ** | 4.26 x 10^-73^ | 6.92 x 10^-8^ | 6.41 x 10^-9^ |

3. **Statistical analysis and post hoc tests for the Specificity**

**SFig.2** illustrates the specificity metric of the kNN classifier using the PID metric applied to brain networks derived from healthy controls and MCI. Results are presented across the frequency bands and the six approaches, based on representative time series, multivariate extensions of bivariate connectivity measures, and multivariate connectivity estimators.

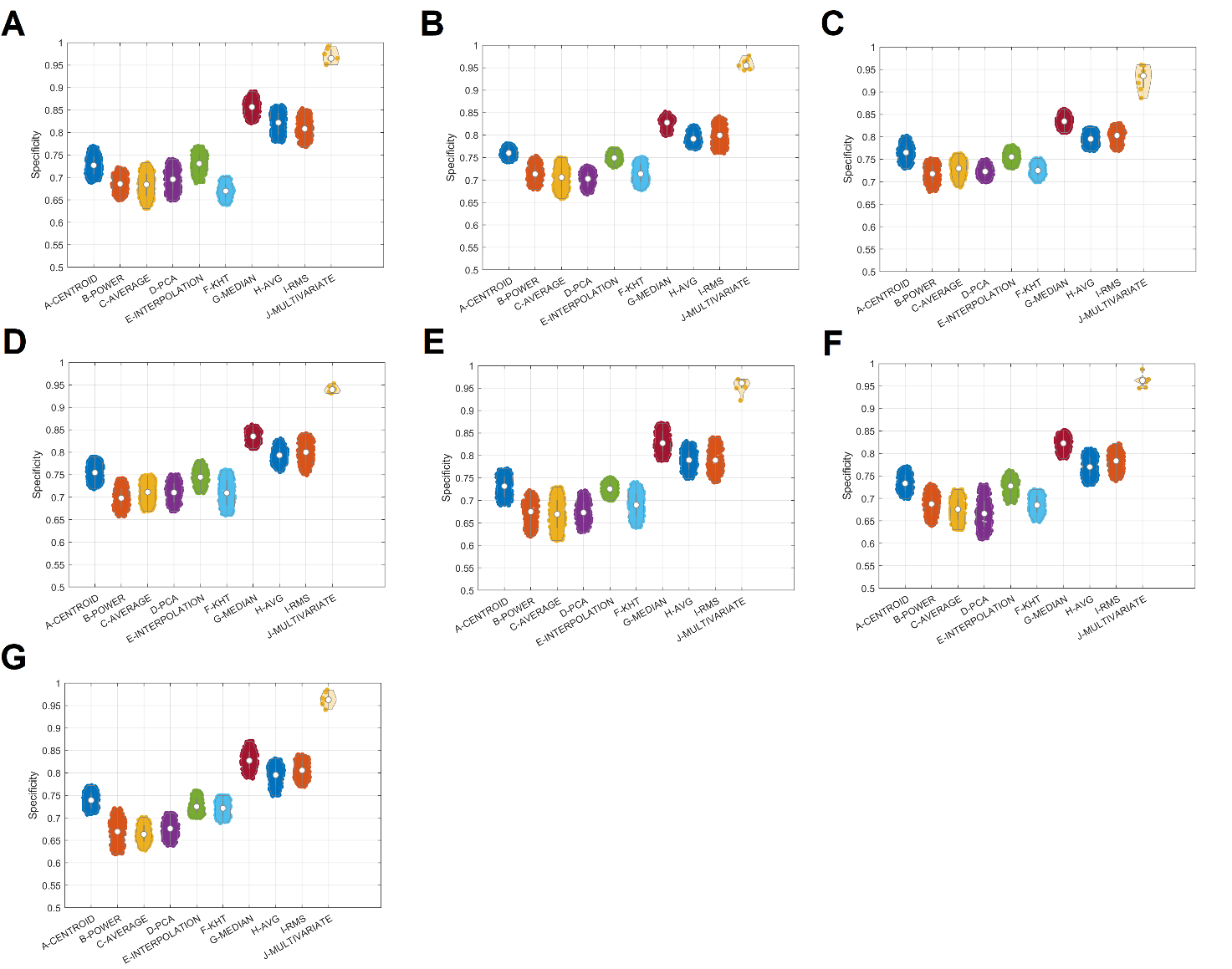

**SFig.2.** Specificity of the machine learning approach applied over individual functional brain networks across the frequency bands, and the different approaches. (A-G : δ,θ,α_1_,α_2_,β_1_,β_2_,γ frequency bands)

**3.1 Representative time series**

**STable 11** summarizes the results of Friedman's test for the six approaches, based on representative time series across the frequency bands. Results from Nemenyi's test and Cliff’s delta for each frequency band are summarized in **STable 12.**

**STable 11.** Results of Friedman’s Test for the six approaches based on representative time series across the frequency bands.

|  | **df** | **χ^2^** | **P-value** |
| --- | --- | --- | --- |
| **δ** | 5 | 591.97 | 1.10 x 10^-125^ |
| **θ** | 5 | 763.45 | 9.33 x 10^-163^ |
| **α_1_** | 5 | 630.95 | 4.15 x 10^-134^ |
| **α_2_** | 5 | 514.59 | 5.66 x 10^-109^ |
| **β_1_** | 5 | 611.63 | 6.21 x 10^-130^ |
| **β_2_** | 5 | 609.45 | 1.84 x 10^-129^ |
| **γ** | 5 | 881.06 | 5.46 x 10^-187^ |

**STable 12. (A – G ).** Summary of Nemenyi's test and Cliff’s delta for each frequency band.

**A.delta frequency**

Method 1	Method 2	LowerLimit	Difference	UpperLimit	Pvalue	Cliffs d

1	2	1.6967	2.1833	2.67	0	0.80195

1	3	1.8008	2.2875	2.7742	0	0.75349

1	4	1.3008	1.7875	2.2742	0	0.61946

1	5	-0.58251	-0.095833	0.39085	0.99345	-0.08145

1	6	2.6508	3.1375	3.6242	0	0.97336

2	3	-0.38251	0.10417	0.59085	0.99036	0.066527

2	4	-0.88251	-0.39583	0.090847	0.18676	-0.17144

2	5	-2.7658	-2.2792	-1.7925	0	-0.8251

2	6	0.46749	0.95417	1.4408	3.4245e-07	0.4228

3	4	-0.98668	-0.5	-0.01332	0.039979	-0.2227

3	5	-2.87	-2.3833	-1.8967	0	-0.78326

3	6	0.36332	0.85	1.3367	9.5587e-06	0.25764

4	5	-2.37	-1.8833	-1.3967	0	-0.66276

4	6	0.86332	1.35	1.8367	2.2264e-14	0.50739

5	6	2.7467	3.2333	3.72	0	0.97047

**B. Theta frequency**

Method 1	Method 2	LowerLimit	Difference	UpperLimit	Pvalue	Cliffs d

1	2	2.3925	2.8792	3.3658	0	0.96524

1	3	2.7592	3.2458	3.7325	0	0.97266

1	4	3.0383	3.525	4.0117	0	1

1	5	0.084153	0.57083	1.0575	0.010756	0.44027

1	6	2.2925	2.7792	3.2658	0	0.95248

2	3	-0.12001	0.36667	0.85335	0.26315	0.19386

2	4	0.15915	0.64583	1.1325	0.0021569	0.32374

2	5	-2.795	-2.3083	-1.8217	0	-0.8636

2	6	-0.58668	-0.1	0.38668	0.99202	-0.047629

3	4	-0.20751	0.27917	0.76585	0.57542	0.07901

3	5	-3.1617	-2.675	-2.1883	0	-0.88916

3	6	-0.95335	-0.46667	0.020013	0.068976	-0.23257

4	5	-3.4408	-2.9542	-2.4675	0	-0.99944

4	6	-1.2325	-0.74583	-0.25915	0.00018268	-0.36084

5	6	1.7217	2.2083	2.695	0	0.82695

C. **Alpha1 frequency**

**Method 1	Method 2	LowerLimit	Difference	UpperLimit	Pvalue	Cliffs d**

1	2	2.7217	3.2083	3.695	0	0.90917

1	3	1.9133	2.4	2.8867	0	0.79522

1	4	2.3175	2.8042	3.2908	0	0.91614

1	5	-0.065847	0.42083	0.90751	0.13486	0.28389

1	6	2.205	2.6917	3.1783	0	0.89226

2	3	-1.295	-0.80833	-0.32165	3.2561e-05	-0.2917

2	4	-0.89085	-0.40417	0.082513	0.16809	-0.17915

2	5	-3.2742	-2.7875	-2.3008	0	-0.86925

2	6	-1.0033	-0.51667	-0.029987	0.029904	-0.24257

3	4	-0.082513	0.40417	0.89085	0.16809	0.14913

3	5	-2.4658	-1.9792	-1.4925	0	-0.70443

3	6	-0.19501	0.29167	0.77835	0.5265	0.087378

4	5	-2.87	-2.3833	-1.8967	0	-0.87918

4	6	-0.59918	-0.1125	0.37418	0.9863	-0.084972

5	6	1.7842	2.2708	2.7575	0	0.84418

D. **Alpha2 frequency**

**Method 1	Method 2	LowerLimit	Difference	UpperLimit	Pvalue	Cliffs d**

1	2	2.5467	3.0333	3.52	0	0.92887

1	3	1.8342	2.3208	2.8075	0	0.80513

1	4	1.8842	2.3708	2.8575	0	0.83288

1	5	-0.053347	0.43333	0.92001	0.11337	0.24296

1	6	1.855	2.3417	2.8283	0	0.75188

2	3	-1.1992	-0.7125	-0.22582	0.00043308	-0.29086

2	4	-1.1492	-0.6625	-0.17582	0.0014662	-0.26851

2	5	-3.0867	-2.6	-2.1133	0	-0.86273

2	6	-1.1783	-0.69167	-0.20499	0.00072787	-0.21614

3	4	-0.43668	0.05	0.53668	0.99971	0.028033

3	5	-2.3742	-1.8875	-1.4008	0	-0.69062

3	6	-0.46585	0.020833	0.50751	1	0.026813

4	5	-2.4242	-1.9375	-1.4508	0	-0.72626

4	6	-0.51585	-0.029167	0.45751	0.99998	0.0062413

5	6	1.4217	1.9083	2.395	0	0.63532

E. **Beta 1 frequency**

**Method 1	Method 2	LowerLimit	Difference	UpperLimit	Pvalue	Cliffs d**

1	2	2.2467	2.7333	3.22	0	0.88013

1	3	2.2425	2.7292	3.2158	0	0.82371

1	4	2.2425	2.7292	3.2158	0	0.86618

1	5	-0.46585	0.020833	0.50751	1	0.1106

1	6	1.3758	1.8625	2.3492	0	0.66879

2	3	-0.49085	-0.0041667	0.48251	1	0.046897

2	4	-0.49085	-0.0041667	0.48251	1	-0.041946

2	5	-3.1992	-2.7125	-2.2258	0	-0.93944

2	6	-1.3575	-0.87083	-0.38415	5.063e-06	-0.34317

3	4	-0.48668	0	0.48668	1	-0.080265

3	5	-3.195	-2.7083	-2.2217	0	-0.85704

3	6	-1.3533	-0.86667	-0.37999	5.7562e-06	-0.33912

4	5	-3.195	-2.7083	-2.2217	0	-0.92211

4	6	-1.3533	-0.86667	-0.37999	5.7562e-06	-0.31862

5	6	1.355	1.8417	2.3283	0	0.69174

F. **Beta 2 frequency**

**Method 1	Method 2	LowerLimit	Difference	UpperLimit	Pvalue	Cliffs d**

1	2	1.98	2.4667	2.9533	0	0.85656

1	3	2.3467	2.8333	3.32	0	0.94055

1	4	2.6758	3.1625	3.6492	0	0.88856

1	5	-0.06168	0.425	0.91168	0.12738	0.23853

1	6	1.9758	2.4625	2.9492	0	0.91217

2	3	-0.12001	0.36667	0.85335	0.26315	0.2446

2	4	0.20915	0.69583	1.1825	0.00065691	0.31555

2	5	-2.5283	-2.0417	-1.555	0	-0.74508

2	6	-0.49085	-0.0041667	0.48251	1	0.010112

3	4	-0.15751	0.32917	0.81585	0.38521	0.12709

3	5	-2.895	-2.4083	-1.9217	0	-0.86008

3	6	-0.85751	-0.37083	0.11585	0.25119	-0.24979

4	5	-3.2242	-2.7375	-2.2508	0	-0.82636

4	6	-1.1867	-0.7	-0.21332	0.00059249	-0.31838

5	6	1.5508	2.0375	2.5242	0	0.79745

G. **Gamma frequency**

**Method 1	Method 2	LowerLimit	Difference	UpperLimit	Pvalue	Cliffs d**

1	2	2.8675	3.3542	3.8408	0	0.98135

1	3	3.1967	3.6833	4.17	0	1

1	4	2.6175	3.1042	3.5908	0	0.99878

1	5	0.16332	0.65	1.1367	0.0019604	0.36318

1	6	0.44665	0.93333	1.42	6.8705e-07	0.51199

2	3	-0.15751	0.32917	0.81585	0.38521	0.086925

2	4	-0.73668	-0.25	0.23668	0.68744	-0.11953

2	5	-3.1908	-2.7042	-2.2175	0	-0.92772

2	6	-2.9075	-2.4208	-1.9342	0	-0.87134

3	4	-1.0658	-0.57917	-0.092487	0.0090944	-0.27381

3	5	-3.52	-3.0333	-2.5467	0	-0.99951

3	6	-3.2367	-2.75	-2.2633	0	-0.97775

4	5	-2.9408	-2.4542	-1.9675	0	-0.96185

4	6	-2.6575	-2.1708	-1.6842	0	-0.89819

5	6	-0.20335	0.28333	0.77001	0.55911	0.2001

3.2 **Multivariate Approaches for bivariate measures**

**STable 13.** summarizes the results of Friedman's test for the three approaches of the multivariate extension of bivariate FC estimators across the frequency bands. Results from Nemenyi's test and Cliff’s delta for each frequency band are summarized in **STable 14.**

**STable 13.** Results of Friedman’s Test for the three approaches of the multivariate extension of bivariate FC estimators across the frequency bands.

|  | **df** | **χ^2^** | **P-value** |
| --- | --- | --- | --- |
| **δ** | 2 | 218.13 | 4.29 x 10^-48^ |
| **θ** | 2 | 188.58 | 1.12 x 10^-41^ |
| **α_1_** | 2 | 310.26 | 4.24 x 10^-68^ |
| **α_2_** | 2 | 259.27 | 5.00 x 10^-57^ |
| **β_1_** | 2 | 176.7 | 4.26 x 10^-39^ |
| **β_2_** | 2 | 291.26 | 5.67 x 10^-64^ |
| **γ** | 2 | 162.61 | 4.89 x 10^-36^ |

**STable 14. (A – G ).** Summary of Nemenyi's test and Cliff’s delta for each frequency band.

**A. Delta – frequency**

Method 1	Method 2	LowerLimit	Difference	UpperLimit	Pvalue	Cliffs d

1	2	0.81938	1.0333	1.2473	0	0.71695

1	3	1.0527	1.2667	1.4806	0	0.85673

2	3	0.019384	0.23333	0.44728	0.02854	0.24501

**B. Theta – frequency**

Method 1	Method 2	LowerLimit	Difference	UpperLimit	Pvalue	Cliffs d

1	2	0.94855	1.1625	1.3764	0	0.85938

1	3	0.77355	0.9875	1.2014	0	0.59386

2	3	-0.38895	-0.175	0.03895	0.13384	-0.11653

**C. Alpha1 – frequency**

Method 1	Method 2	LowerLimit	Difference	UpperLimit	Pvalue	Cliffs d

1	2	1.3027	1.5167	1.7306	0	0.95314

1	3	1.0069	1.2208	1.4348	0	0.86722

2	3	-0.50978	-0.29583	-0.081884	0.0034169	-0.22413

D. **Alpha 2 – frequency**

Method 1	Method 2	LowerLimit	Difference	UpperLimit	Pvalue	Cliffs d

1	2	1.1611	1.375	1.5889	0	0.91824

1	3	0.92355	1.1375	1.3514	0	0.75589

2	3	-0.45145	-0.2375	-0.02355	0.025146	-0.14059

**E. Beta 1 – frequency**

Method 1	Method 2	LowerLimit	Difference	UpperLimit	Pvalue	Cliffs d

1	2	0.86105	1.075	1.2889	0	0.72897

1	3	0.81105	1.025	1.2389	0	0.66698

2	3	-0.26395	-0.05	0.16395	0.84764	-0.01834

**F. Beta 2 – frequency**

Method 1	Method 2	LowerLimit	Difference	UpperLimit	Pvalue	Cliffs d

1	2	1.2694	1.4833	1.6973	0	0.919

1	3	0.94022	1.1542	1.3681	0	0.80987

2	3	-0.54312	-0.32917	-0.11522	0.00090788	-0.27043

**G.Gamma frequency**

Method 1	Method 2	LowerLimit	Difference	UpperLimit	Pvalue	Cliffs d

1	2	0.94438	1.1583	1.3723	0	0.70492

1	3	0.46522	0.67917	0.89312	2.6311e-13	0.51607

2	3	-0.69312	-0.47917	-0.26522	4.5767e-07	-0.32019

We applied the Aligned Rank Transform (ART) ANOVA, a nonparametric factorial method equivalent to 2-way ANOVA, to assess the potential effects of frequency bands, the six approaches for estimating representative time series, the three approaches for the multivariate extension of bivariate estimators, and their interactions on classification performance. Our findings found only an effect of the methods and no effect of the frequency bands and their interactions (df = 5, F = 1129.84, p = 5.38 x 10^-17^, effect of the six approaches for estimating the representative time series, and df = 2, F = 987.62, p = 4.12 x 10^-15^, effect of the three approaches for the multivariate extension of the bivariate connectivity estimators).

3.3 **The classification performance of Multivariate estimators vs the approaches of representative time series and the multivariate extension of bivariate connectivity measures**

**STable 15** summarizes p-values derived from the application of the Wilcoxon rank sum test between pairs of methodologies and across the frequency bands for the sensitivity measurements.

**STable 15.** Summary of p-values derived from the application of the Wilcoxon rank sum test between pairs of methodologies and across the frequency bands for the sensitivity measurements. 1 refers to approaches for estimating representative time series, 2 to the multivariate extension of bivariate connectivity measures, and 3 to multivariate estimators.

|  | **1 vs 2** | **1 vs 3** | **2 vs 3** |
| --- | --- | --- | --- |
| **δ** | 5.42 x 10^-79^ | 4.42 x 10^-7^ | 5.44 x 10^-6^ |
| **θ** | 3.72 x 10^-74^ | 6.72 x 10^-7^ | 6.23 x 10^-7^ |
| **α_1_** | 3.39 x 10^-77^ | 5.38 x 10^-6^ | 1.38 x 10^-6^ |
| **α_2_** | 5.64 x 10^-78^ | 6.25 x 10^-6^ | 5.83 x 10^-5^ |
| **β_1_** | 4.29 x 10^-76^ | 5.24 x 10^-5^ | 2.36 x 10^-5^ |
| **β_2_** | 3.05 x 10^-74^ | 3.82 x 10^-5^ | 5.71 x 10^-7^ |
| **γ** | 4.72 x 10^-75^ | 5.27 x 10^-8^ | 5.98 x 10^-6^ |

4. **Statistical analysis and post hoc tests for the Repeatability Cohort**

**SFig.3** illustrates the specificity metric of the kNN classifier using the PID metric applied to brain networks derived from healthy controls and MCI. Results are presented across the frequency bands and the six approaches, based on representative time series, multivariate extensions of bivariate connectivity measures, and multivariate connectivity estimators.

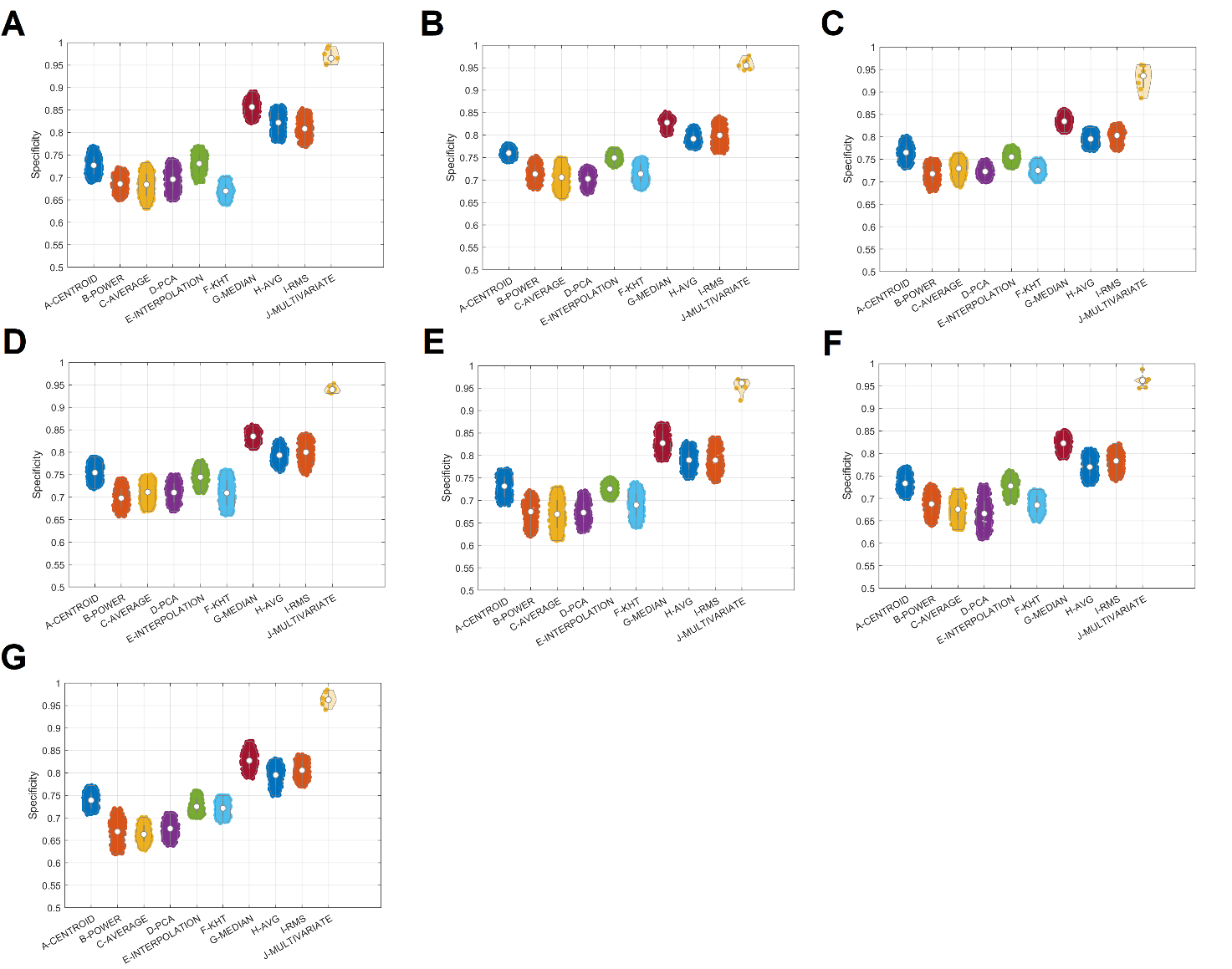

**SFig.3.** Repeatability of the functional brain network topologies measured with PID and computed between sessions across the frequency bands, and the different approaches. (A-G : δ,θ,α_1_,α_2_,β_1_,β_2_,γ frequency bands)

**4.1 Representative time series**

**STable 16** summarizes the results of Friedman's test for the six approaches, based on representative time series across the frequency bands. Results from Nemenyi's test and Cliff’s delta for each frequency band are summarized in **STable 17.**

**STable 16.** Results of Friedman’s Test for the six approaches based on representative time series across the frequency bands.

|  | **df** | **χ^2^** | **P-value** |
| --- | --- | --- | --- |
| **δ** | 5 | 482.3 | 5.26 x 10^-102^ |
| **θ** | 5 | 772.27 | 1.44 x 10^-164^ |
| **α_1_** | 5 | 645.45 | 3.04 x 10^-137^ |
| **α_2_** | 5 | 486.82 | 5.58 x 10^-103^ |
| **β_1_** | 5 | 517.67 | 1.22 x 10^-109^ |
| **β_2_** | 5 | 594.66 | 2.87 x 10^-126^ |
| **γ** | 5 | 480.95 | 1.03 x 10^-101^ |

**STable 17. (A – G ).** Summary of Nemenyi's test and Cliff’s delta for each frequency band.

**A.delta frequency**

**Method 1**	**Method 2**	**LowerLimit**	**Difference**	**UpperLimit**	**Pvalue**	**Cliffs d**
 1	2	-2.8533	-2.3667	-1.88	0	0
 1	3	-2.6158	-2.1292	-1.6425	0	0.26667
 1	4	-2.7575	-2.2708	-1.7842	0	-0.066667
 1	5	-0.34501	0.14167	0.62835	0.96213	0.26667
 1	6	-2.5867	-2.1	-1.6133	0	-0.066667
 2	3	-0.24918	0.2375	0.72418	0.73277	0.066667
 2	4	-0.39085	0.095833	0.58251	0.99345	-0.13333
 2	5	2.0217	2.5083	2.995	0	0.2
 2	6	-0.22001	0.26667	0.75335	0.62413	0
 3	4	-0.62835	-0.14167	0.34501	0.96213	-0.33333
 3	5	1.7842	2.2708	2.7575	0	0.066667
 3	6	-0.45751	0.029167	0.51585	0.99998	-0.066667
 4	5	1.9258	2.4125	2.8992	0	0.26667
 4	6	-0.31585	0.17083	0.65751	0.918	0
 5	6	-2.7283	-2.2417	-1.755	0	-0.26667

**B. Theta frequency**

**Method 1**	**Method 2**	**LowerLimit**	**Difference**	**UpperLimit**	**Pvalue**	**Cliffs d**
 1	2	-3.1867	-2.7	-2.2133	0	-0.2
 1	3	-3.7492	-3.2625	-2.7758	0	-0.26667
 1	4	-3.4367	-2.95	-2.4633	0	-0.13333
 1	5	-0.75751	-0.27083	0.21585	0.60796	-0.26667
 1	6	-3.6283	-3.1417	-2.655	0	0
 2	3	-1.0492	-0.5625	-0.07582	0.012685	0
 2	4	-0.73668	-0.25	0.23668	0.68744	0.066667
 2	5	1.9425	2.4292	2.9158	0	0
 2	6	-0.92835	-0.44167	0.045013	0.10059	0.46667
 3	4	-0.17418	0.3125	0.79918	0.44638	0.26667
 3	5	2.505	2.9917	3.4783	0	0.066667
 3	6	-0.36585	0.12083	0.60751	0.98111	0.4
 4	5	2.1925	2.6792	3.1658	0	-0.26667
 4	6	-0.67835	-0.19167	0.29501	0.87233	0.2
 5	6	-3.3575	-2.8708	-2.3842	0	0.33333

**C.Alpha1 frequency**

**Method 1**	**Method 2**	**LowerLimit**	**Difference**	**UpperLimit**	**Pvalue**	**Cliffs d**
 1	2	-2.9658	-2.4792	-1.9925	0	0.2
 1	3	-3.2492	-2.7625	-2.2758	0	0.26667
 1	4	-3.1533	-2.6667	-2.18	0	0.46667
 1	5	-0.69501	-0.20833	0.27835	0.82732	0.26667
 1	6	-3.52	-3.0333	-2.5467	0	0.2
 2	3	-0.77001	-0.28333	0.20335	0.55911	0.2
 2	4	-0.67418	-0.1875	0.29918	0.88243	0.46667
 2	5	1.7842	2.2708	2.7575	0	-0.13333
 2	6	-1.0408	-0.55417	-0.067487	0.01492	0.066667
 3	4	-0.39085	0.095833	0.58251	0.99345	0.46667
 3	5	2.0675	2.5542	3.0408	0	-0.33333
 3	6	-0.75751	-0.27083	0.21585	0.60796	0
 4	5	1.9717	2.4583	2.945	0	-0.53333
 4	6	-0.85335	-0.36667	0.12001	0.26315	-0.33333
 5	6	-3.3117	-2.825	-2.3383	0	0.066667

D. **Alpha2 frequency**

**Method 1**	**Method 2**	**LowerLimit**	**Difference**	**UpperLimit**	**Pvalue**	**Cliffs d**
 1	2	-2.6908	-2.2042	-1.7175	0	-0.13333
 1	3	-2.9158	-2.4292	-1.9425	0	-0.2
 1	4	-2.62	-2.1333	-1.6467	0	-0.066667
 1	5	-0.52001	-0.033333	0.45335	0.99996	0.33333
 1	6	-2.9617	-2.475	-1.9883	0	-0.33333
 2	3	-0.71168	-0.225	0.26168	0.77548	-0.066667
 2	4	-0.41585	0.070833	0.55751	0.99844	-0.066667
 2	5	1.6842	2.1708	2.6575	0	0.26667
 2	6	-0.75751	-0.27083	0.21585	0.60796	-0.2
 3	4	-0.19085	0.29583	0.78251	0.51027	0.13333
 3	5	1.9092	2.3958	2.8825	0	0.4
 3	6	-0.53251	-0.045833	0.44085	0.99981	-0.066667
 4	5	1.6133	2.1	2.5867	0	0.33333
 4	6	-0.82835	-0.34167	0.14501	0.34184	-0.26667
 5	6	-2.9283	-2.4417	-1.955	0	-0.53333

E. **Beta 1 frequency**

**Method 1	Method 2	LowerLimit	Difference	UpperLimit	Pvalue	Cliffs d**

1	2	-2.3575	-1.8708	-1.3842	0	0.26667
 1	3	-2.6908	-2.2042	-1.7175	0	-0.13333
 1	4	-2.295	-1.8083	-1.3217	0	0.2
 1	5	0.12582	0.6125	1.0992	0.0045243	-0.2
 1	6	-2.6908	-2.2042	-1.7175	0	0.066667
 2	3	-0.82001	-0.33333	0.15335	0.37048	-0.4
 2	4	-0.42418	0.0625	0.54918	0.99915	-0.066667
 2	5	1.9967	2.4833	2.97	0	-0.26667
 2	6	-0.82001	-0.33333	0.15335	0.37048	0.066667
 3	4	-0.090847	0.39583	0.88251	0.18676	0.26667
 3	5	2.33	2.8167	3.3033	0	-0.066667
 3	6	-0.48668	0	0.48668	1	0.33333
 4	5	1.9342	2.4208	2.9075	0	-0.26667
 4	6	-0.88251	-0.39583	0.090847	0.18676	-0.13333
 5	6	-3.3033	-2.8167	-2.33	0	0.33333

F. **Beta 2 frequency**

**Method 1	Method 2	LowerLimit	Difference	UpperLimit	Pvalue	Cliffs d**

1	2	-2.87	-2.3833	-1.8967	0	0
 1	3	-3.2325	-2.7458	-2.2592	0	0.066667
 1	4	-2.9408	-2.4542	-1.9675	0	0.066667
 1	5	-0.57418	-0.0875	0.39918	0.99572	-0.066667
 1	6	-3.2158	-2.7292	-2.2425	0	0.066667
 2	3	-0.84918	-0.3625	0.12418	0.27545	0
 2	4	-0.55751	-0.070833	0.41585	0.99844	-0.066667
 2	5	1.8092	2.2958	2.7825	0	-0.13333
 2	6	-0.83251	-0.34583	0.14085	0.32794	0.066667
 3	4	-0.19501	0.29167	0.77835	0.5265	-0.066667
 3	5	2.1717	2.6583	3.145	0	-0.066667
 3	6	-0.47001	0.016667	0.50335	1	0.066667
 4	5	1.88	2.3667	2.8533	0	-0.13333
 4	6	-0.76168	-0.275	0.21168	0.59172	0.066667
 5	6	-3.1283	-2.6417	-2.155	0	0.066667

G. **Gamma frequency**

**Method 1	Method 2	LowerLimit	Difference	UpperLimit	Pvalue	Cliffs d**
 1	2	-2.6408	-2.1542	-1.6675	0	0.33333
 1	3	-2.6617	-2.175	-1.6883	0	-0.13333
 1	4	-2.9575	-2.4708	-1.9842	0	-0.066667
 1	5	-0.52001	-0.033333	0.45335	0.99996	0.13333
 1	6	-2.8783	-2.3917	-1.905	0	0.4
 2	3	-0.50751	-0.020833	0.46585	1	-0.46667
 2	4	-0.80335	-0.31667	0.17001	0.43078	-0.33333
 2	5	1.6342	2.1208	2.6075	0	-0.066667
 2	6	-0.72418	-0.2375	0.24918	0.73277	0
 3	4	-0.78251	-0.29583	0.19085	0.51027	0.13333
 3	5	1.655	2.1417	2.6283	0	0.26667
 3	6	-0.70335	-0.21667	0.27001	0.8022	0.53333
 4	5	1.9508	2.4375	2.9242	0	0.2
 4	6	-0.40751	0.079167	0.56585	0.99734	0.33333
 5	6	-2.845	-2.3583	-1.8717	0	0.13333

4.2 **Multivariate Approaches for bivariate measures**

**STable 18.** summarizes the results of Friedman's test for the three approaches of the multivariate extension of bivariate FC estimators across the frequency bands. Results from Nemenyi's test and Cliff’s delta for each frequency band are summarized in **STable 19.**

**STable 18.** Results of Friedman’s Test for the three approaches of the multivariate extension of bivariate FC estimators across the frequency bands.

|  | **df** | **χ^2^** | **P-value** |
| --- | --- | --- | --- |
| **δ** | 2 | 29.78 | 3.42 x 10^-7^ |
| **θ** | 2 | 13.41 | 0.0012 |
| **α_1_** | 2 | 15.83 | 0.0004 |
| **α_2_** | 2 | 4.41 | 0.11 |
| **β_1_** | 2 | 7.68 | 0.02 |
| **β_2_** | 2 | 35.76 | 1.71 x 10^-8^ |
| **γ** | 2 | 9.86 | 0.0072 |

**STable 19. (A – G ).** Summary of Nemenyi's test and Cliff’s delta for each frequency band.

**A. Delta – frequency**

Method 1	Method 2	LowerLimit	Difference	UpperLimit	Pvalue	Cliffs d

1	2	-0.20145	0.0125	0.22645	0.98972	-0.33333
 1	3	-0.63895	-0.425	-0.21105	9.6372e-06	-0.33333
 2	3	-0.65145	-0.4375	-0.22355	4.9185e-06	0

**B. Theta – frequency**

Method 1	Method 2	LowerLimit	Difference	UpperLimit	Pvalue	Cliffs d

1	2	0.10688	0.32083	0.53478	0.0012804	-1
 1	3	0.027717	0.24167	0.45562	0.022108	-0.66667
 2	3	-0.29312	-0.079167	0.13478	0.66101	0.33333

**C. Alpha1 – frequency**

Method 1	Method 2	LowerLimit	Difference	UpperLimit	Pvalue	Cliffs d

1	2	-0.54728	-0.33333	-0.11938	0.00076211	0.33333
 1	3	-0.25562	-0.041667	0.17228	0.89154	1
 2	3	0.077717	0.29167	0.50562	0.0039949	0.66667

D. **Alpha 2 – frequency**

Method 1	Method 2	LowerLimit	Difference	UpperLimit	Pvalue	Cliffs d

1	2	1.1611	1.375	1.5889	0	0.91824

1	3	0.92355	1.1375	1.3514	0	0.75589

2	3	-0.45145	-0.2375	-0.02355	0.025146	-0.14059

**E. Beta 1 – frequency**

Method 1	Method 2	LowerLimit	Difference	UpperLimit	Pvalue	Cliffs d

1	2	-0.22645	-0.0125	0.20145	0.98972	0
 1	3	-0.43895	-0.225	-0.01105	0.036528	0
 2	3	-0.42645	-0.2125	0.0014496	0.052043	-0.33333

**F. Beta 2 – frequency**

Method 1	Method 2	LowerLimit	Difference	UpperLimit	Pvalue	Cliffs d

1	2	-0.71812	-0.50417	-0.29022	9.9807e-08	-0.33333
 1	3	-0.64728	-0.43333	-0.21938	6.1672e-06	-0.33333
 2	3	-0.14312	0.070833	0.28478	0.71783	0

**G.Gamma frequency**

Method 1	Method 2	LowerLimit	Difference	UpperLimit	Pvalue	Cliffs d

1	2	-0.43062	-0.21667	-0.0027171	0.046351	0.33333
 1	3	-0.48478	-0.27083	-0.056884	0.0084564	0
 2	3	-0.26812	-0.054167	0.15978	0.82368	-0.33333

We applied the Aligned Rank Transform (ART) ANOVA, a nonparametric factorial method equivalent to 2-way ANOVA, to assess the potential effects of frequency bands, the six approaches for estimating representative time series, the three approaches for the multivariate extension of bivariate estimators, and their interactions on repeatability. Our findings found only an effect of the methods and no effect of the frequency bands and their interactions (df = 5, F = 1466.57, p = 2.22 x 10^-308^, effect of the six approaches for estimating the representative time series, and df = 2, F = 241.22, p = 2.22 x 10^-308^, effect of the three approaches for the multivariate extension of the bivariate connectivity estimators).

4.3 **The classification performance of Multivariate estimators vs the approaches of representative time series and the multivariate extension of bivariate connectivity measures**

**STable 20** summarizes p-values derived from the application of the Wilcoxon rank sum test between pairs of methodologies and across the frequency bands for the repeatability measurements.

**STable 20.** Summary of p-values derived from the application of the Wilcoxon rank sum test between pairs of methodologies and across the frequency bands for the repeatability measurements. 1 refers to approaches for estimating representative time series, 2 to the multivariate extension of bivariate connectivity measures, and 3 to multivariate estimators.

|  | **1 vs 2** | **1 vs 3** | **2 vs 3** |
| --- | --- | --- | --- |
| **δ** | 4.12 x 10^-80^ | 6.62 x 10^-6^ | 6.62 x 10^-6^ |
| **θ** | 3.64 x 10^-72^ | 5.83 x 10^-7^ | 6.67 x 10^-6^ |
| **α_1_** | 3.82 x 10^-71^ | 5.05 x 10^-5^ | 1.89 x 10^-7^ |
| **α_2_** | 4.34 x 10^-75^ | 6.72 x 10^-7^ | 5.24 x 10^-6^ |
| **β_1_** | 5.37 x 10^-70^ | 4.53 x 10^-6^ | 3.77 x 10^-5^ |
| **β_2_** | 3.67 x 10^-72^ | 4.12 x 10^-5^ | 4.92 x 10^-5^ |
| **γ** | 4.24 x 10^-71^ | 5.62 x 10^-8^ | 5.14 x 10^-6^ |
